# Salinisation and warming disrupt predator-induced drift behaviour in aquatic predator-prey interactions

**DOI:** 10.1101/2024.07.02.601583

**Authors:** Anna-Maria Vermiert, Iris Madge Pimentel, Philipp M. Rehsen, Tobias Otto, Martin Horstmann, Arne J. Beermann, Florian Leese, Linda C. Weiss, Ralph Tollrian

## Abstract

Predators act as a major selective force impacting biological communities in riverine ecosystems. Often predation risk is mediated by chemical cues upon which prey species display distinct morphological or behavioural defences. For stream invertebrates, predator-induced behaviours can be drifting and hiding. However, anthropogenic stressors increasingly impact freshwater ecosystems, with predicted consequences on the chemical information transfer, potentially affecting anti-predator behaviour. Despite this, knowledge and research on the extent to which these stressors may affect evolved predator-induced behaviour and subsequent ecosystems is scarce. Therefore, we conducted an outdoor mesocosm experiment (*ExStream* system) at a lowland stream in Germany, in which we investigated the effects of direct and indirect fish predation on invertebrate communities under different stressor conditions. As stressors, we tested elevated salinity (ambient vs + 136 mg/L NaCl), elevated temperature (ambient vs. + 3.4°C), and the combination of both. Our findings reveal distinct predator-induced drift behaviour, characterised by a greater number of drifting invertebrates and a concurrent reduction in overall invertebrate numbers in the communities. Increased salinity and temperature hampered the predator-induced drift behaviour. Our results support the possibility that stressors negatively impact the prey’s ability to perceive predators’ presence and respond appropriately. This disruption may alter predator-prey relationships, thereby affecting communities and potentially ecosystem functions.

**Highlights:**

▯ We hypothesised that anthropogenic stressors disrupt predator-induced behaviours in invertebrate prey species.
▯ Predator-induced drift behaviour was experimentally studied under ambient and elevated salinity and temperature conditions.
▯ Fish exposure elicited predator-induced drift behaviour, which decreased the number of invertebrate individuals of the communities.
▯ Elevated salt and elevated temperatures alone and in combination did not induce drift behaviour.
▯ We found that both stressors disrupted predator-induced drift behaviour, altering the anti-predator response of invertebrate prey.

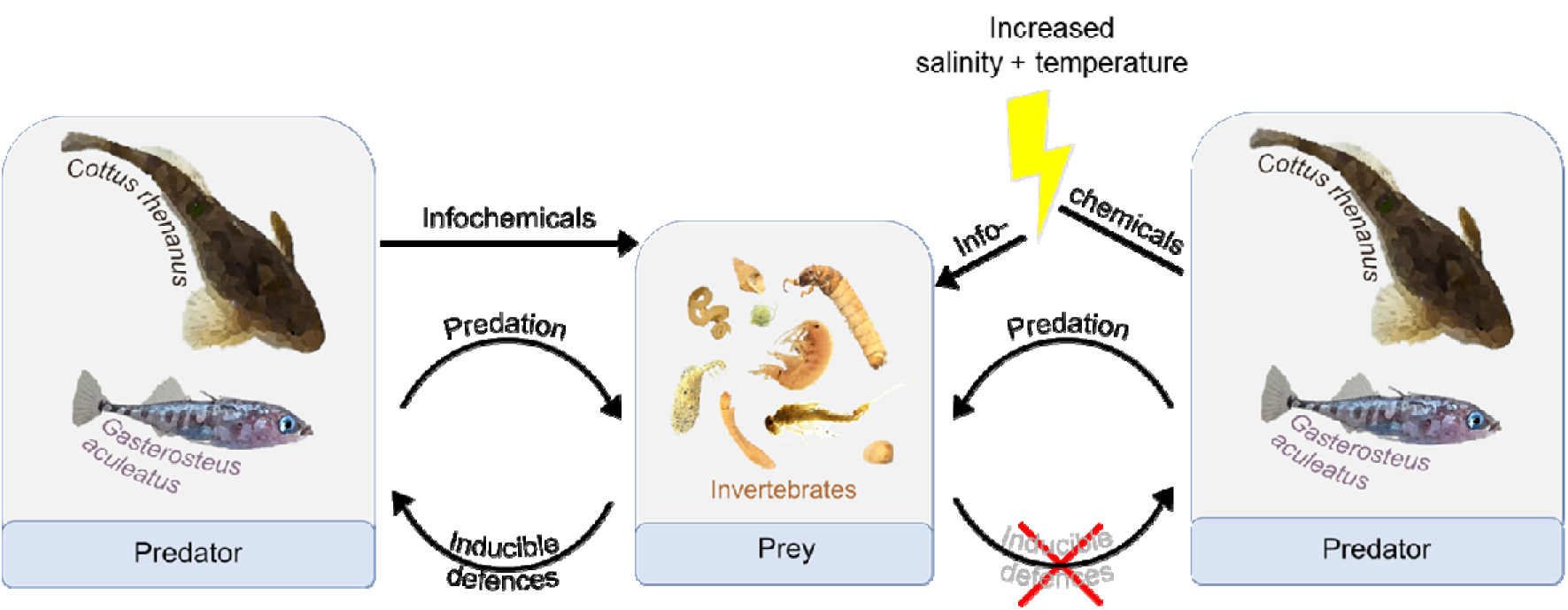

## 1 Introduction

Predation shapes the behaviour, morphology, and distribution in ecosystems worldwide. From terrestrial to aquatic environments, prey species have evolved an extensive array of defence mechanisms and avoidance strategies in response to predation (Caro, 2014; Dettner, 2019). In lotic freshwater ecosystems, predatory fish exert strong pressure on various invertebrates (Crowder and Cooper, 1982; Dahl, 1998a; Williams et al., 2003). Invertebrate taxa are known to exhibit distinct predator avoidance strategies (Dettner, 2019; Kasumyan, 2022; Weiss and Tollrian, 2018). A common strategy of numerous invertebrate taxa is drifting, characterised as a passive transport of aquatic organisms with the water current (Müller, 1954; Waters, 1972). This movement with the current downstream can also be used to evade unfavourable conditions (Dahl, 1998b; Poff et al., 1991). Other prevalent strategies observed in invertebrates are the downregulation of activity levels to reduce visibility and predator-induced hiding behaviour, where the organisms seek refuge in, e.g., sediment or vegetation (Ahlgren et al., 2011; Lauridsen and Lodge, 1996). Such anti-predator strategies are often first induced upon the perception of chemical cues, such as alarm cues from conspecifics and/or predatory cues (Gall and Brodie, 2009; Huryn and Chivers, 1999; Weiss, 2019). Many prey species have an evolved sensitivity to these cues which consequently act as an early warning system to assess the predation risk and respond appropriately (Ferrari et al., 2010; Helfman, 1989; Huryn and Chivers, 1999).

Recent studies show that several anthropogenic stressors interfere with predator-induced responses and can modify or disrupt organismic interactions (Trotter et al., 2019; Weiss et al., 2018). At sublethal levels, they could, for example, either affect the release or persistence of chemical cues in the aquatic environment, or the stressors change the mode of signal transmission or perception. As an example, elevated levels of carbon dioxide inhibit predator cue recognition in freshwater habitats, as demonstrated for salmon and *Daphnia* (Ou et al., 2015; Weiss et al., 2018). Similarly, Van Donk et al. (2015) reasoned that even low concentrations of drugs interfere with natural chemical information transfer due to their structural and functional similarities to infochemicals.

Salinisation and warming are two globally prevalent stressors on riverine ecosystems. Freshwater salinisation can be caused, for instance, by agricultural irrigation, industrial sources, and the use of road de-icers (Cañedo-Argüelles et al., 2019; Shtull-Trauring et al., 2020). It can have varying effects on a broad range of freshwater invertebrates depending on their sensitivity, causing both lethal and non-lethal impacts (Castillo et al., 2018; Hart et al., 1991). This is in part due to the direct impacts of salt ions on osmoregulation, ultimately reducing cellular stability, and leading to higher energetic costs to maintain homeostasis (Buchwalter et al., 2019; Cañedo-Argüelles et al., 2019). When the salinity level exceeds the tolerance threshold of an organism, the effect results in higher mortality rates (Berezina, 2003; Willems et al., 2022). Moreover, some studies have demonstrated that elevated salinity levels can disrupt conspecific communication in, e.g., fish (*Pseudomugil signifer*) and lobster (*Panulirus argus*) (Herbert-Read et al., 2010; Ross and Behringer, 2019) and influences the perception of alarm or predatory cues (Hintz and Relyea, 2017; Huber et al., 2023; Jones et al., 2017; Reustle and Smee, 2020). Therefore, higher salinity may also influence intracellular ion distribution, with consequences for chemoreceptor neurons, their response to neuronal stimuli, and the patterns of activity they exhibit in response to these stimuli (Ross and Behringer, 2019).

In comparison to increased salinity, increased temperature often has more contrasting effects on organisms. Effects can be directly beneficial if the temperature increase is towards an organism’s optimal thermal range, or negative if the thermal tolerance range is exceeded (Dallas and Ketley, 2011; Dallas and Ross-Gillespie, 2015; Vannote and Sweeney, 1980). Furthermore, indirect effects of increased temperature must also be considered, e.g., reduced oxygen levels in warmer water with at the same time higher metabolic rates. In the context of climate change, but also due to the removal of riparian vegetation, stream ecosystems face increasing temperatures (Kail et al., 2021; Woodward et al., 2010). The effects of increased temperature that could alter predator-induced behaviour and predator recognition include, e.g., increased metabolic rates and thus increased activity of predators and prey. This metabolic increase can result in increased cue production, leading to a higher concentration of chemical cues (Draper and Weissburg, 2019). Additionally, heightened activity due to increased metabolism may render predators and prey more visible to each other as movement increases. Increased temperature can also affect chemical reactions, including catabolism. Higher temperatures can accelerate the degradation of cues through multiple mechanisms (e.g., increased molecular motion, enhanced enzyme activity, accelerated evaporation, oxidative reaction), thereby reducing their concentration in the environment (Draper and Weissburg, 2019). This potential for temperature stress to cause both an increase and decrease in chemical cue concentration leaves the specific mechanism undetermined, but it likely depends on the magnitude of the temperature difference. In both cases, the change in concentration could change the reaction of prey to predation.

Individually and in combination, the effects of salinization and warming could therefore affect the predator-induced responses, potentially hampering anti-predator behaviour. However, the effects of these stressors on organismic interactions are not well understood. We therefore investigated their effects on predator-induced prey species behaviour and its impact on the invertebrate community.

We hypothesised that: 1.) Predators induce drift behaviour in prey. 2) Predator-induced drift behaviour will reduce the overall number of invertebrates remaining in the community. 3.) Salinization and warming, both individually and in combination, disrupt predator-induced behaviour.

To test our hypotheses, we conducted an outdoor mesocosm experiment using the two stressors – increased salinity (+136 mg/L NaCl) and elevated temperature (+ 3.4 °C) - with and without predation exposure in a full factorial design. To evaluate the impact on predator-induced behaviour, we quantified the proportion of invertebrates that evaded the system via drift and those that hid in the leaf litter under stress conditions. Additionally, we compared the invertebrate numbers in the leaf litter and substrate under these conditions, assessing how predator-induced behaviour influenced the invertebrate community.

## 2 Material and methods

### 2.1 Study site

The Boye catchment is situated in the Ruhr region in North Rhine-Westphalia, Germany, and spans 77 square kilometres. The Boye river is a carbonate-rich, sand-bottom lowland river (www.elwasweb.nrw.de; OFWK ID: DE_NRW_27726, last accessed June 20, 2024). Since the beginning of the 20th century, it has been used as an open sewer for wastewater from mining, industry, domestic sewage, and agriculture. In 2017, as part of the Generation Project Emscher Restoration, a separated wastewater channel was constructed and since then the Boye river has undergone a restoration towards a near-natural state. Prior to restoration, only Oligochaeta inhabited sections of the Boye stream (Winking et al., 2016). Since then, a rich macroinvertebrate community has reestablished, fuelled by intact tributary sections in the catchment area (Gillmann et al., 2023). The restoration of the Boye section within the study area was completed in 2021. Our electrofishing conducted by Dr C. Edler and Birgit Daniel (Bezirksregierung Düsseldorf, Dezernat 51: Nature and landscape protection, fishery) revealed a substantial population of sticklebacks (Gasterosteidae), a few bullheads (Cottidae), and occasionally other fish species such as bleak and sunbleak (Cyprinidae).

### 2.2 Experimental design

On March 4, 2022, the outdoor mesocosm system (*ExStream* system; Piggott et al., 2015) was established adjacent to the Boye river (An der Boy, Gladbeck; decimal degrees: N 51.5533°, E 6.9485°). The *ExStream* system allows for the investigation of different stressor effects on a stream ecosystem by providing a high number of replications and maintaining precise control over experimental variables. At the same time, it has a high degree of realism due to its capability of achieving identical physico-chemical conditions as the adjacent stream and permitting invertebrate migration (Beermann et al., 2018; Piggott et al., 2015; Wagenhoff et al., 2012).

The system consisted of 64 identical flow-through channels and was based on a scaffolding with a top and lower level (Fig. 1). Two pumps (Pedrollo NGAm 1A-Pro) transported the Boye water to the top of the scaffolding system. To shield the pumps from larger debris and prevent fish from entering the system, the pump’s intake hoses were encased in protective covers with 5 mm diameter holes. At the top level, four sets of three 203 L collection tanks connected in series were placed (Fig. 2). The two rear tanks functioned as sediment traps and were installed to avert extreme sedimentation and clogging of the system. The water passively flowed through the sediment traps into the front tank designated as the header tank (Fig. 2). Each header tank was connected to 16 mesocosms via hoses, allowing water to flow gravity-driven to the mesocosms at the lower level (Fig. 1, Fig. 2). The flow rate of the mesocosm channel was maintained at approximately 2.1 L/min via a flow valve. To reflect the substrates of the Boye streambed, the mesocosms were filled with 1000 mL of sediment collected from the Boye catchment (N 51.5544°, E 6.9463°; sieved through a 1 mm sieve), 100 mL of fine particulate organic matter (FPOM, N 51.5627°, E 6.9154°), 200 mL of quartz materials (6-8 mm), and 3 larger stones (40-80 mm). In addition to sediment, a single coarse-meshed leaf litter bag, filled with approximately 7 g of alder leaf litter (*Alnus glutinosa*), measuring 2.5 cm in diameter and 15 cm in length, with a mesh size of 5 mm, was inserted into each of the mesocosms (Fig. 1B). Leaves were collected during the previous autumn and then air-dried at room temperature.

**Fig. 1:**
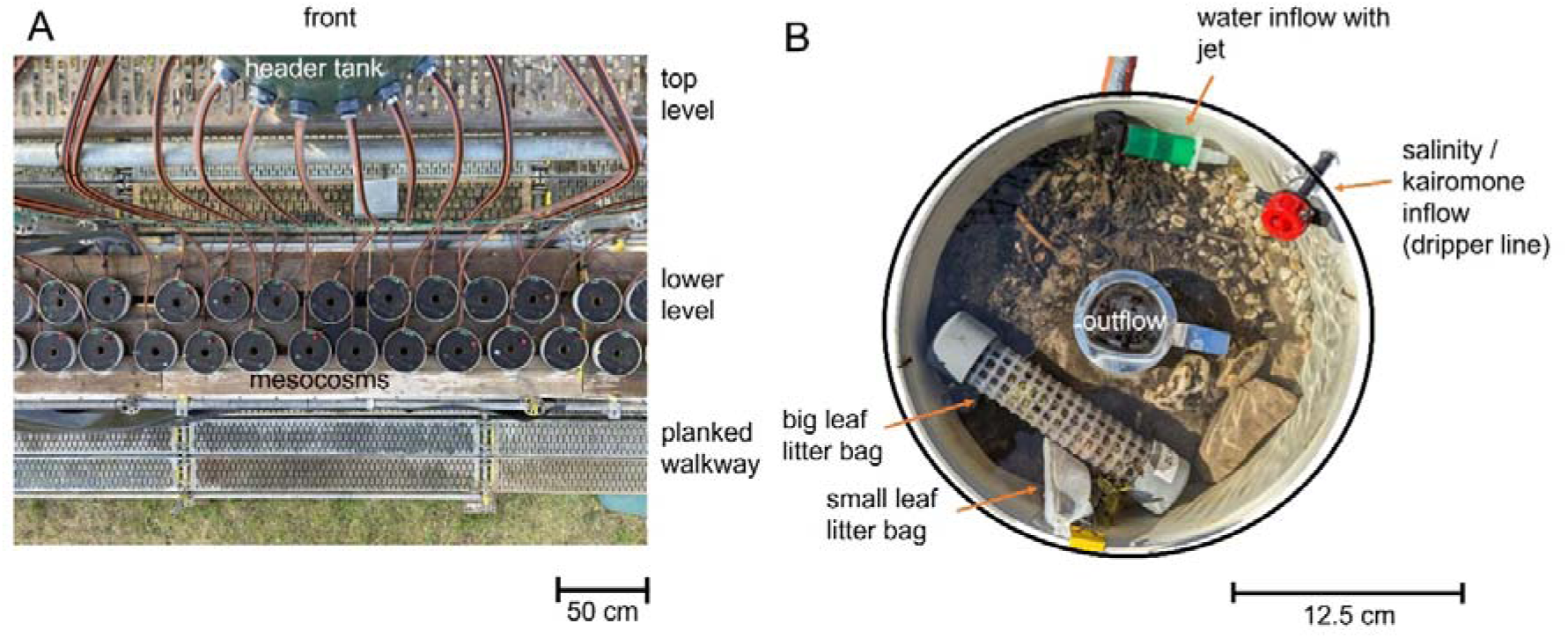
Setup of the *ExStream* system and single mesocosm. (A) Top-down view of the experimental *ExStream* system setup with level definition. (B) Circular mesocosm (volume 3.5 L; Microwave Ring Moulds, Interworld, Auckland, New Zealand) was filled with streambed substrate and two leaf litter bags. Water in the mesocosm had a clockwise flow direction and drained through the central outflow. To study predator-induced drift behaviour, nets were placed in the outflow to capture the invertebrates leaving the system. Invertebrates remaining in the leaf litter bags were observed for predator-induced hiding behaviour. The invertebrates remaining in the system, including those in the leaf litter bag, sediment and water column, were considered part of the community. They were used to investigate the impact of predator exposure, stressors and both typical and altered predator-avoidance behaviour on the invertebrate community.

**Fig. 2:**
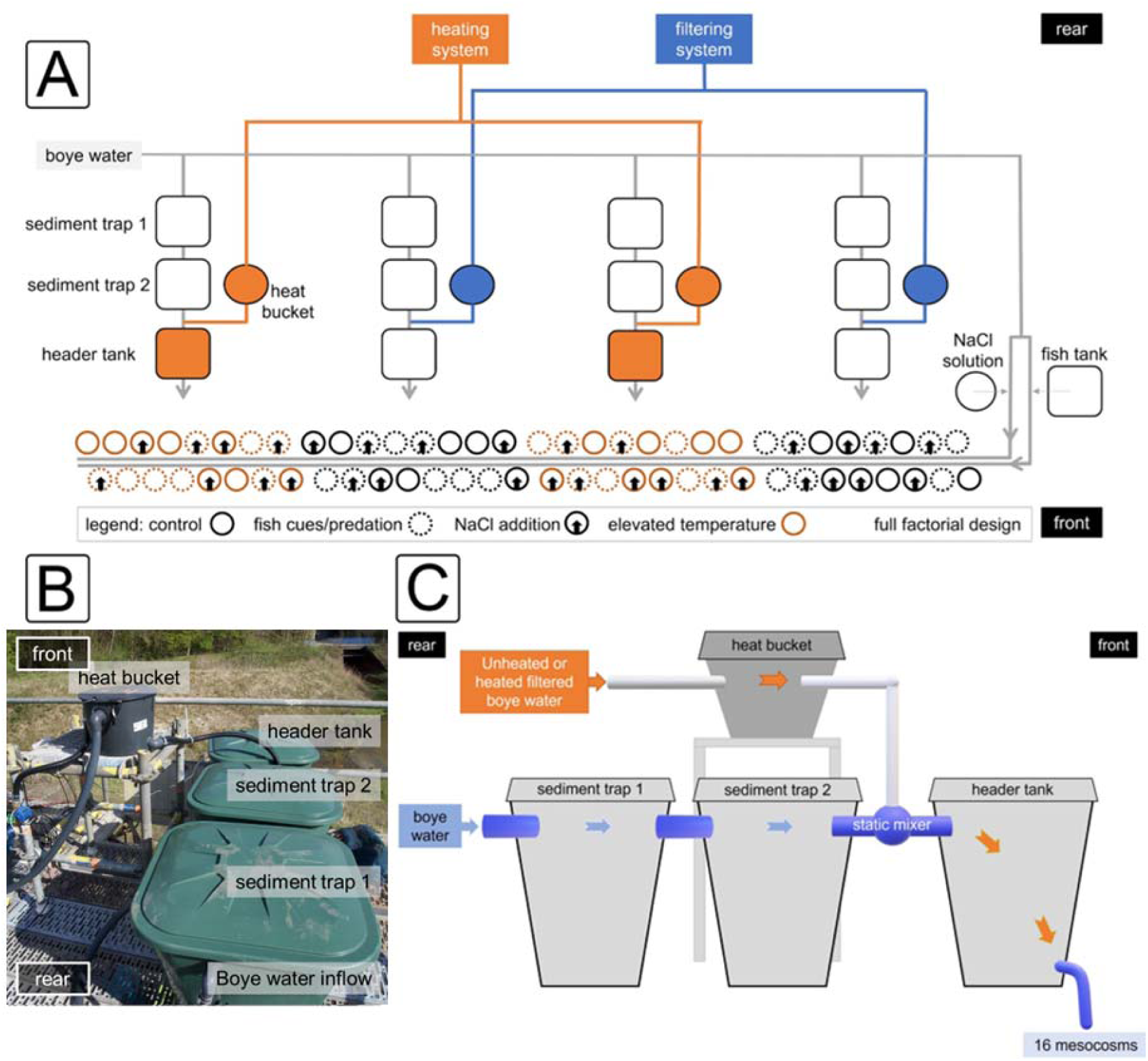
Experimental set-up of the *ExStream* system (A) Schematic overview of the experimental mesocosm system with 64 mesocosms assigned in four randomized blocks. Stressors in the mesocosms are indicated by dotted circles (predator exposure), black arrows (increased salinity) and orange circles (increased temperature). The direction of water flow is indicated by grey arrows. Fish tank water and NaCl solution were applied via two separate irrigation lines to the mesocosms. (B) Photo and (C) Schematic overview of the sediment traps, the header tank and the heat bucket with the connections at the top level. The water flow is indicated by blue and when heated by orange arrows. The sediment trap and header tank are located on the top level of the scaffold above the mesocosms (lower level). The connection of the second sediment trap and header tank is linked to the heating system. Heated water, at approximately 50 °C, is stored in a bucket and fed into the system once the water temperature of the header tank is less than 4 °C above the water temperature of the sediment trap.

The *ExStream* system was set-up in a full factorial design to examine the impact of increased salinity and temperature on direct and/or indirect predator exposure. The salinity stressor and predator exposure treatment were randomly assigned in 4 blocks, each with 16 mesocosms, while increased temperature was implemented per block (Fig 2A). The water that passed through the system was directed to a retention basin afterwards.

#### 2.2.1 Predator exposure

This study was conducted in accordance with the Animal Welfare Act. To implement predator exposure, two fish species - three-spined sticklebacks (Gasterosteidae: *Gasterosteus aculeatus*) and bullheads (Cottidae: *Cottus rhenanus*) - were captured and distributed between two tanks (internal dimensions 113 cm x 93 cm x 57.5 cm) set up on ground level. For each fish tank, an additional tank served as a water reservoir to maintain the water level by means of a mechanical pump. The sticklebacks, ranging from 4 to 7 cm in size, were captured via electrofishing at the study site. Bullheads, with a size range of 3.8 to 7.2 cm (mean: 5.72, SD: 0.78), were caught through night fishing in the Wannebach (N 51.4245°, E 8.0511°), which is a tributary of the Ruhr River located in the urban area of Arnsberg. To emulate natural feeding behaviour and environmental conditions, we provided the caught fish with a daily portion of invertebrates gathered from a downstream section.

During predator exposure implementation, invertebrates in the mesocosms were initially exposed to fish-enriched water (indirect predation) and subsequently to the direct presence of a bullhead (direct predation). For indirect predation, the sticklebacks and bullheads were distributed evenly between two fish tanks. The estimated total biomass of stickleback and bullhead in the tanks was 480 g. Using a peristaltic pump (Hei-FLOW Value 01 with a multi-channel pump head C8) the water from each tank was pumped over two separate hoses (Tygon hoses: standard R3603, inner diameter 6.4 mm, wall thickness 1.6 mm, outer diameter 9.5 mm) into the same irrigation line. The fish-enriched water (0.62 g/L equalling 100 % of predator cues) was pumped at a constant rate of 32.4 L/h through the irrigation line connected to a pressure-compensated dripper system (4 L/h per mesocosm). Diluted by the Boye water flowing through the irrigation line and the mesocosms, the fish-treated mesocosms had a final fish concentration of 0.005 g/L (0.8% of the predator cues, indirect predation). Due to a technical failure, the peristaltic pump stopped working for one night during the stressor phase (day 12).

Additionally, to the 0.005 g/L fish-enriched water, direct predator exposure was implemented on the fifth day of the stressor phase. For this, as a second stimulation phase and to test for the consumptive effect of direct predation, bullheads (*Cottus rhenanus*) were placed directly in the mesocosms. Hence, the bullheads (31) were collected from the fish tanks to place one of them directly into each predator-treated mesocosm (indirect and direct predator exposure). Bullheads have been successfully kept in a similar setup before (Matthaei personal communication). Unexpectedly, 14 bullheads escaped out of our mesocosms within the first six days and eight more till the end of the stressor phase. It is possible that the proximity to the mating season between March and April (Kottelat and Freyhof, 2007) was too close and an important aspect of their activity was searching for a mating partner. Thus, an estimation of the consumptive effect of predation was not possible. Yet, the direct introduction of fish resulted in a notable increase in fish biomass concentration in the mesocosms, reaching around 16 mg/L. The concentration increase, combined with other cues elicited by the short presence of the bullhead, facilitated a second opportunity to test for predator-induced drift behaviour. At the end of the experiment, all escaped fish were collected in an outflow basin. The bullheads and sticklebacks were returned to the area where they were collected and released.

#### 2.2.2 Elevated salinity levels

The stressor salinity was implemented via a NaCl solution. The NaCl solution (refined salt tablets, Claramat, > 99.9 % NaCl, >350 g/L NaCl) was collected in a 1 m^3^ plastic tank next to the setup. It was then pumped over the salt irrigation line into the mesocosms using a pressure-compensated dripper system (see Beermann et al., 2018). Since the dripper line was constantly supplied with filtered Boye water, the NaCl solution entering the mesocosms was diluted (136 mg/L NaCl). This increased the ambient conductivity (i.e., ambient salinity) from an average of 840.63 µS/cm ± 94.20 SD (median 853 µS/cm) to an average of 1446.06 µS/cm ± 912.39 SD (median 1296 µS/cm). The salinity level was chosen based on the maximum conductivity observed across the Boye system. After an initial runtime of a few hours, the salt application was discontinued due to technical failure by one day. The salt stressor was reactivated at the beginning of the second stressor day. In addition, deactivation of the salt occurred for 12 hours during the night from the 12^th^ to the 13^th^ day of the stressor phase due to flooding of the river. Several issues related to clogging also resulted in higher conductivity than anticipated (Supplementary fig. 1).

#### 2.2.3 Elevated temperature levels

The ambient temperature of the mesocosm water averaged 8.7 °C, with temperatures as low as 2.3 °C or up to 16.1 °C degrees occurring during the stressor phase. The target temperature in the header tank was set to + 4.5 °C (± 0.2 °C) relative to the ambient water temperature in the sediment trap to account for heat loss as the water travels to the mesocosms (targeted + 4.0 °C). The difference in water temperature between warmed and ambient mesocosms averaged 3.4°C (± 0.8°C). To ensure that the bullheads were not exposed to temperatures above their thermal limits, the maximum temperature was set to 24 °C (Elliott and Elliott, 1995). Heating was achieved through the heat exchange mechanism of a petroleum-driven heater with an internal water circuit, preceded by a filtration system. The filtered Boye water was heated to approximately 50 °C, pumped up onto the scaffold, and redirected to heat buckets (Fig. 2). The outflow of these buckets was controlled via temperature sensors in the sediment trap 2 and header tank with a magnetic valve, and continuously increased the water temperature of the header tank to + 4.5 °C (± 0.2 °C) (Madge Pimentel et al., 2024b). The two blocks with ambient water temperature were constructed identically to those with heated water, except they received filtered water via a second filtration system not connected to the heating unit (Fig. 2A). The outflow of buckets with filtered water was kept identical to the outflow from the adjacent heat buckets to ensure equal amounts of filtered water in the header tanks of the temperature and ambient treatments. Due to high sediment loads in the stream’s water after strong precipitation events, the temperature stressor was turned off temporarily on several occasions on days 5, 6, 8, 13 and 14 of the stressor phase (Supplementary fig. 2).

### 2.3 Experimental procedures

The experiment was divided into two consecutive phases: A 20-day colonisation phase and a 14-day stressor phase (Fig. 3). During the colonisation phase natural communities were established. Coarse-meshed leaf litter tubes were inserted into the mesocosm on the fourth day of the colonisation phase. They provided shelter and additional food for invertebrates and were used to study predator-induced hiding behaviour. As natural immigration of invertebrates with a diameter larger than 5 mm was prevented due to the protective cover around the intake hose, the communities were supplemented by multi-habitat kick-net sampling from an upstream part of the Boye on the fifth day of the colonisation phase. These samples were collected from around 51 streambed patches. The macroinvertebrates caught were added in similar densities to the experimental channels using a mixing procedure developed by Elbrecht et al. (2016). The gathered macroinvertebrates were collected in a tank filled with Boye water, stirred thoroughly and 5 litres were scooped out. Using a bucket, 8 identical jars were arranged in a circle and placed on a potter’s wheel. While the wheel rotated, the previously scooped liquid was poured out evenly till the jars were filled. The content from two filled jars was added to one mesocosm. This was repeated until all the mesocosms received the water and macroinvertebrates of one jar.

**Fig. 3:**
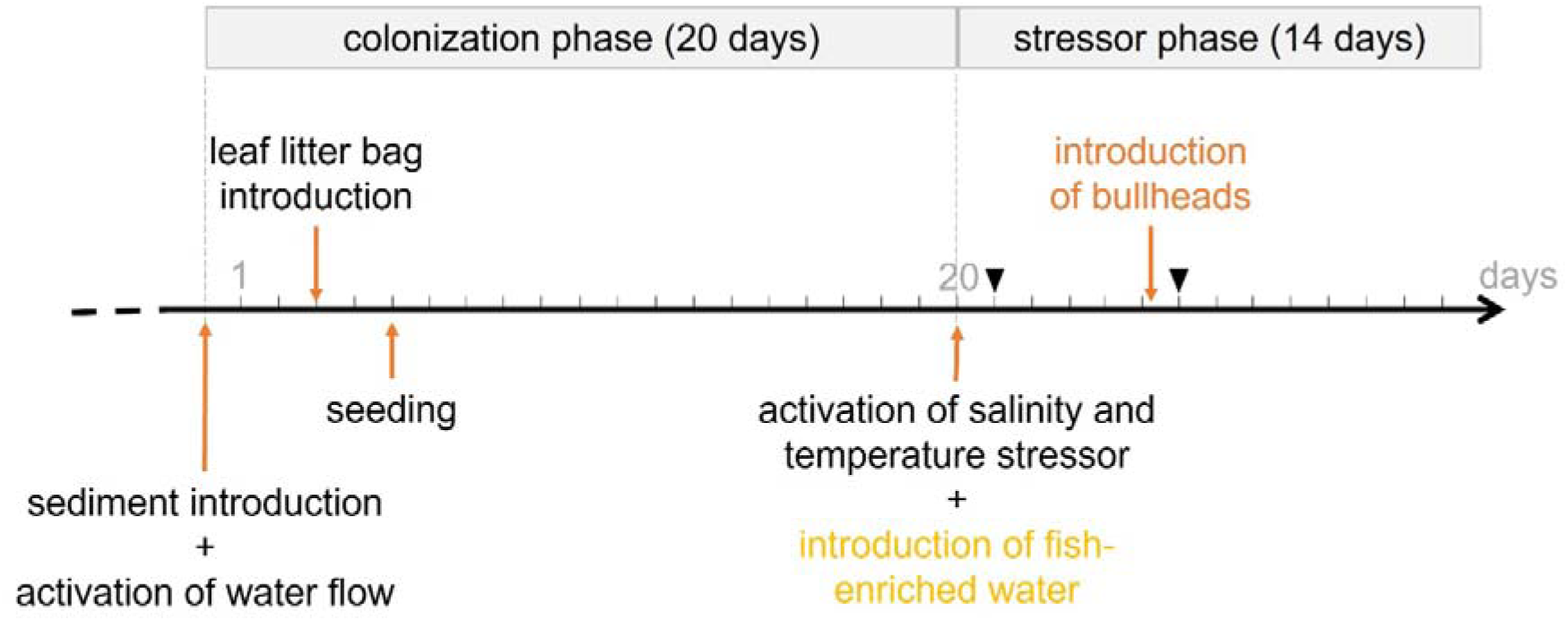
Schedule for the *ExStream* system. After setup, the 20-day colonization phase began with sediment being added to the mesocosms and water flow being activated. This was followed by leaf litter placement and a seeding event that introduced additional macroinvertebrates. The 14-day stressor phase began with the introduction of salinity and temperature stressors and the introduction of fish-enriched water (indirect predation). At midnight of the fifth day, bullheads were placed in the mesocosms (direct predation). The two drift sample days are marked with a black triangle.

To manage the high sediment loads in the tanks, nets (mesh size 10 mm) were hung in the second sediment traps four days before the start of the stressor phase to reduce floating sediment particles from reaching the header tank. Additionally, the tanks were cleared of sediment multiple times using a pool cleaner. During this process, the flow in the mesocosms was briefly stopped in the mesocosms using a valve, and the initial water load of the mesocosm inflow was diverted outside the mesocosm to prevent stirred-up sediment from entering.

With the start of the stressor phase, the two stressors, i.e., increased salinity and increased temperature, and predator exposure were applied in a full-factorial design. To study normal and altered predator-induced drift behaviour, nylon nets were affixed to the outflow for 24 hours, capturing any drifting organisms attempting to exit the system via water current. Drift samples were collected on two days during the stressor phase (day 1 and day 6) to investigate potential behavioural differences of invertebrates influenced by indirect or direct predation. While drift samples were collected from all 64 mesocosms in the experiment, only half of the mesocosms (32) were used as endpoints after 14 days of stressor input. The other half of the mesocosms were maintained for an additional two weeks in a subsequent recovery experiment independent of this study. As a result, only 32 of the leaf litter bags and the mesocosm contents including substrate (substrate sample), were collected.

### 2.4 Sample collection and preparation

The collected drift nets (drift samples) were preserved in 96% ethanol. The invertebrates were later detached from the fabric under running water and collected on a 160 µm sieve. The invertebrates in the leaf litter bags were separated from the leaf litter and preserved in 96% ethanol. Invertebrate samples were subsequently processed via a 250 µm sieve. In the case of substrate samples, the larger stones were first removed and the contents of the mesocosms were filtered through a 500 µm sieve and stored in 96 % ethanol. Smaller stones and sediment were separated from organic matter by elutriation.

### 2.5 Data analysis

Invertebrates were identified using the criteria of Beermann et al. (2018). Using a microscope and standard keys (Bauernfeind and Humpesch, 2001; Eiseler, 2010; Eiseler and Hess, 2015; Lubini et al., 2012; Schaefer, 2018; Sundermann and Lohse, 2004; Waringer and Graf, 1997), invertebrates were identified to the lowest practicable taxonomic level based on morphological characterisation. Rare taxa were grouped into higher taxonomic levels to ensure statistically analysable sample sizes. Additionally, nematodes as well as insects that were too damaged or too small for identification, were excluded from the calculations. Oligochaetes were also excluded from the calculations of the drift data, as they only consisted of small torn-off pieces under 2 mm. To determine the number of oligochaetes in the leaf litter and substrate samples, we distinguished between whole and torn individuals. We then halved the number of broken fragments before adding them to the total count of whole oligochaetes.

The data analysis was performed with R version 4.1.1 (R Core Team, 2021). The effects of the factors predator exposure, salinity, and temperature, as well as their interactions, were analysed for all count data including drift, leaf litter, and community samples (leaf litter + substrate). Community samples were won by aggregating the leaf litter and substrate data as we regarded all invertebrates that remained in the system as part of the community. The number of invertebrates that remained in the system was used to evaluate the effects of predator-induced behaviour, both with and without being altered by stressor input, on the stream community. The invertebrates from the substrate were not analysed separately.

Deviation coding was applied to all categorical variables. The count data were fitted using a generalised linear model (stats package; R Core Team (2021), glm(abundance ∼ fish x salinity x temperature, family = poisson (link = “identity”), data = data)). Given the factorial design of our study, we utilised the identity link function and the Poisson family to address non-normality, while still accounting for interactions on the additive scale. Afterwards, we checked (package AER; Kleiber and Zeileis (2008), dispersiontest(model)) and corrected the Poisson regression models for overdispersion by adjusting the standard error with the estimated dispersion parameter with a quasi-GLM. For all samples, we analysed the total number of all invertebrates and of different taxa/groups depending on the sample using a single model for each. Using post hoc comparisons among groups, conditional effects were calculated when a tendential or significant interaction was found (package emmeans; Lenth (2023)). We corrected the significance threshold of these results for multiple testing using the false discovery rate (FDR).

To analyse the effects of indirect (day 1) and direct predation (day 6) on predator-induced drift behaviour, the mentioned generalised linear model was adapted with the factor day: glm(abundance ∼ day + fish x salinity x temperature, family = poisson (link = “identity”), data = data) The dataset consisted of n = 64 drift data points at day 1 and after the removal of outliers n = 61 drift data points at day 6. One of the mesocosms was excluded from the day 6 data set as it was identified as an extreme outlier, with values exceeding twice the standard deviation. Additionally, two more mesocosms were omitted due to elevated salinity levels surpassing 15,000 µS/cm, which was attributed to water flow obstruction. The drift data were analysed with a focus on specific groups, including chironomid larvae, microcrustaceans (Copepoda, Anomopoda, Ostracoda, and Nauplius larvae), gammarids (*Gammarus* sp.), and EPT taxa (mayfly, stonefly and caddisfly larvae). These groups collectively represented more than 90% of the drifting individuals. Chironomidae pupae and imagines were excluded from the analysis, as they could not have displayed predator-induced drift behaviour. Pupae, being generally immobile, and imagines, having likely drowned, were carried away by the current and landed in the outflow.

For leaf litter and community samples (leaf litter + substrate), we initially had a sample size of 32, as only half of the 64 samples served as endpoints for the stressor phase. The sample size was further reduced to 29, with three mesocosms removed from the analysis: two due to exceptionally high conductivity levels at one-time point and one due to a treatment difference — it lacked a bullhead due to low fish numbers.

In the invertebrate leaf litter and community datasets, we analysed the following taxa that comprised over 90% of the collected invertebrates: Chironomidae larvae (excl. Tanytarsini), Chironomidae pupae, Tanytarsini larvae, Ceratopogonidae larvae, Cyclopoida, Harpacticoida, gammarids, small Trichoptera, Trichoptera over 3 mm and Oligochaeta. The chironomid larvae were separated into Chironmidae larvae, pupae and Tanytarsini larvae in the leaf litter and substrate samples owing to their high abundance. Chironomidae pupae were analysed independently to gather data on potential temporal changes in emergence initiation. Since we primarily observed Trichoptera of small size in the drift, we opted to categorise the Trichoptera into two size groups for the leaf litter and community samples. We separated small Trichoptera measuring between 1 - 3 mm and larger Trichoptera measuring > 3 mm. The Ceratopogonidae larvae were only analysed for the community data as their numbers in leaf litter were too low.

Drift and community results are presented as boxplots bisected at the median value with whiskers reaching values within the 1.5 interquartile range using ggplot (package tidyverse; Wickham et al., 2019).

## 3 Results

### 3.1 Drift behaviour

We found a total of 6753 invertebrate individuals leaving the system via drift (day 1: 4364, day 6: 2389, day 14 remaining total community: 29213). Microcrustaceans (Copepoda, Anomopoda, Ostracoda, and Nauplius larvae) constituted the largest percentage of drifting invertebrates, followed by chironomid larvae, gammarids, and EPT taxa, with other taxa represented in smaller percentages (Fig. 4).

**Fig. 4:**
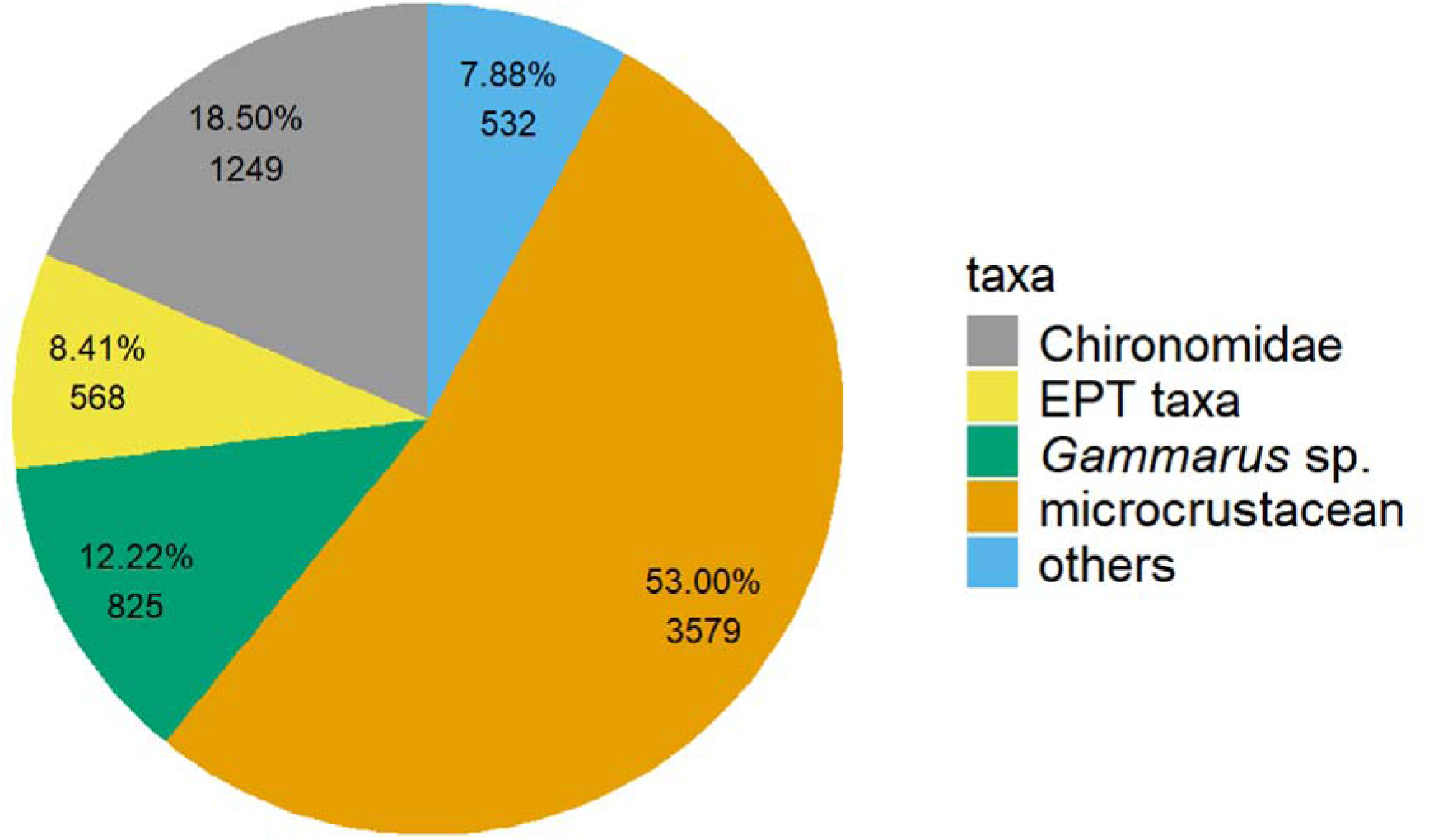
Percentage and number of total drifting invertebrates on two days (day 1, day 6) in a stream mesocosm experiment (*ExStream* system), n = 125.

The number of drifting invertebrates differed between days, with fewer drifting invertebrates observed on the sixth day (glm: df 112, t-value 8.457, Pr(>|t|) < 0.001; Tab. 1). The drift behaviour of all invertebrates was influenced by a three-way interaction of the predator treatment, salinity and temperature (glm: df Inf, t-value -4.197, Pr(>|t|) < 0.001; Tab. 1). Drift in response to predation exposure was exclusively observed under control conditions of the other two stressors (emmeans, indirect predation: df Inf, z-ratio -2.604, p-value 0.014; direct predation: df Inf, z-ratio -3.885, p-value < 0.001; Supplementary tab. 1), or when both temperature and salinity were elevated. When exposed to combined stressors drift was exclusively increased by direct predation, but not by indirect predation (emmeans direct predation: df Inf, z-ratio -4.167, p-value < 0.001; Fig. 5, Supplementary tab. 1). Individually, increased salinity or temperature did not influence the drift behaviour.

**Fig. 5:**
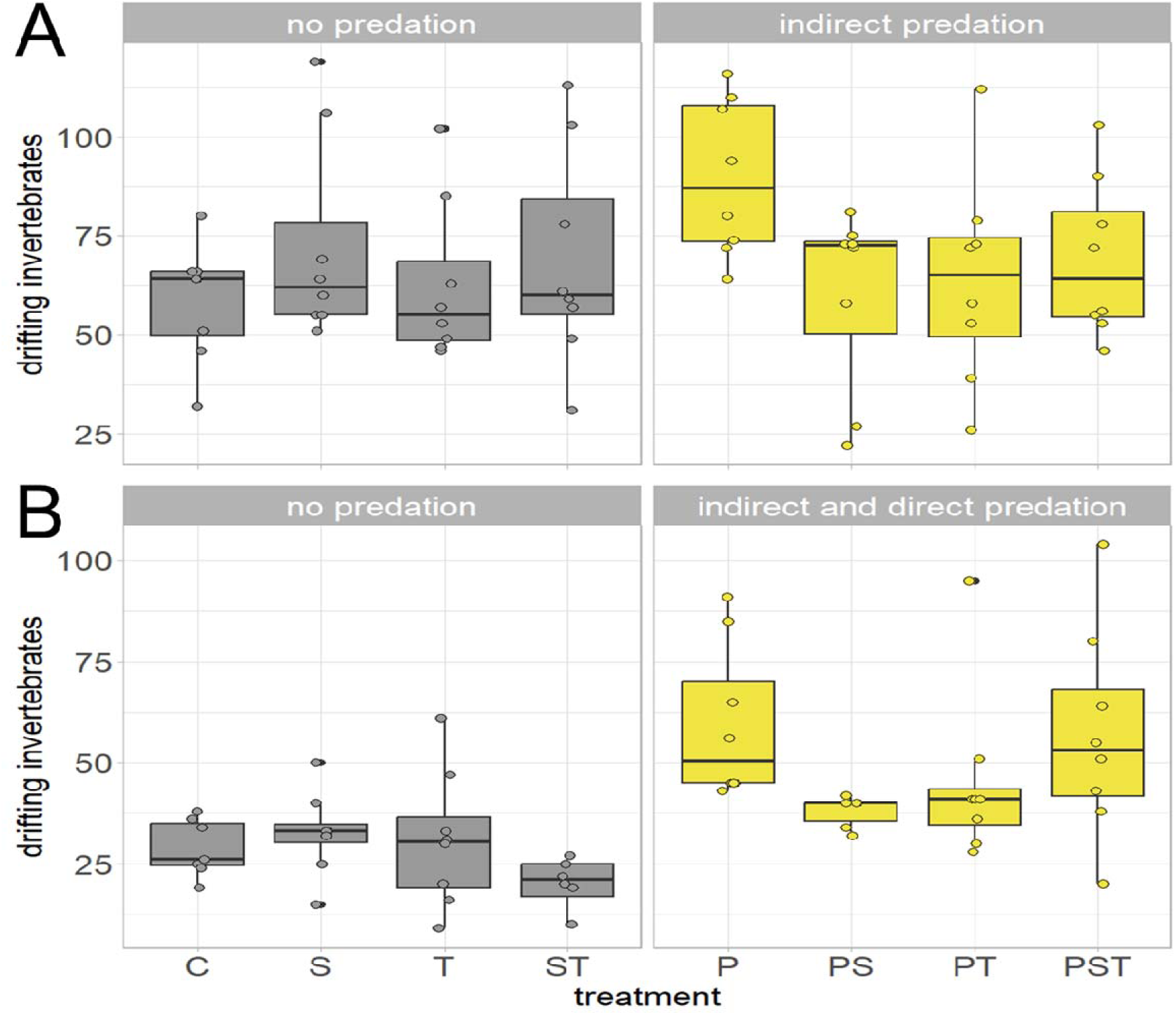
Drifting behaviour under control conditions (C) predator exposure (P) and the stressors increased salinity (S) and increased temperature (T). Total number of drifting invertebrates without (grey) and with predator exposure (yellow) in the field mesocosm experiment on day 1 (A) and day 6 (B). Fish-enriched water (*Gasterosteus aculeatus*, *Cottus rhenanus*) was added continuously from the beginning of the stressor phase (indirect predation). At midnight of the fifth day, bullheads (*Cottus rhenanus*) were added into the mesocosms (direct predation). The data had an n = 64 per day but three outliers were excluded, two of which displayed exceptionally high salinity levels, while the third exhibited twice the standard deviation within the control treatment. Both stressors were divided into two levels, ambient and increased.

**Tab. 1:** Summary of GLM (generalised linear model) results of the invertebrate drifting behaviour of the first and sixth stressor day. Given are the effects and interactions of the factors indirect predator exposure (PI), direct predator exposure (PD), salinity (s), and temperature (t). Predator-specific cues were introduced continuously into the system for indirect predator exposure. On the fifth day, predator exposure was augmented by introducing bullheads into the mesocosms. Bold and italic font denotes tendential (< 0.05) and bold font denotes significant (< 0.1) p-values.

| drift response to predation |  |  |  |  |  |  |  |  |  |  |  |  |  |  |
| --- | --- | --- | --- | --- | --- | --- | --- | --- | --- | --- | --- | --- | --- | --- |
| response variable | values | (Intercept) | day | PD | PI | S | T | PD:S | PI:S | PD:T | PI:T | S:T | PD:S:T | PI:S:T |
| overall | Estimate | 55.990 | 18.236 | -9.474 | -2.478 | 2.506 | 1.890 | -2.721 | 2.558 | 0.013 | 3.296 | 4.251 | -7.544 | 3.376 |
|  | Std. Error | 1.887 | 2.156 | 2.258 | 3.671 | 1.856 | 1.855 | 2.210 | 2.890 | 2.213 | 2.892 | 1.855 | 2.211 | 2.891 |
|  | t value | 29.670 | 8.457 | -4.197 | -0.675 | 1.351 | 1.019 | -1.231 | 0.885 | 0.006 | 1.140 | 2.292 | -3.411 | 1.168 |
|  | Pr(>\|t\|) | **< 0.001** | **< 0.001** | **< 0.001** | 0.501 | 0.180 | 0.311 | 0.221 | 0.378 | 0.996 | 0.257 | **0.024** | **0.001** | 0.245 |
| chironomid larvae | Estimate | 10.250 | 4.742 | -1.405 | -0.787 | 0.705 | 0.807 | -1.247 | 1.527 | -0.104 | 1.149 | 0.693 | -1.708 | 1.164 |
|  | Std. Error | 0.472 | 0.536 | 0.557 | 0.923 | 0.455 | 0.455 | 0.536 | 0.732 | 0.537 | 0.733 | 0.455 | 0.537 | 0.733 |
|  | t value | 21.724 | 8.846 | -2.521 | -0.852 | 1.549 | 1.775 | -2.328 | 2.085 | -0.194 | 1.567 | 1.524 | -3.183 | 1.588 |
|  | Pr(>\|t\|) | **< 0.001** | **< 0.001** | **0.013** | 0.396 | 0.124 | **0.079** | **0.022** | **0.039** | 0.847 | 0.120 | 0.130 | **0.002** | 0.115 |
| microcrustacean | Estimate | 29.682 | 8.438 | -5.243 | 0.143 | 1.708 | 0.739 | -0.851 | 0.153 | 0.121 | 1.775 | 2.560 | -4.523 | 2.302 |
|  | Std. Error | 1.162 | 1.328 | 1.394 | 2.263 | 1.148 | 1.147 | 1.371 | 1.789 | 1.373 | 1.790 | 1.147 | 1.372 | 1.790 |
|  | t value | 25.544 | 6.356 | -3.762 | 0.063 | 1.489 | 0.645 | -0.621 | 0.086 | 0.088 | 0.991 | 2.232 | -3.297 | 1.286 |
|  | Pr(>\|t\|) | **< 0.001** | **< 0.001** | **< 0.001** | 0.950 | 0.139 | 0.521 | 0.536 | 0.932 | 0.930 | 0.324 | **0.028** | **0.001** | 0.201 |
| gammarids | Estimate | 7.061 | 2.142 | -1.794 | -1.318 | 0.422 | 0.412 | -0.428 | 0.694 | -0.029 | 0.472 | 0.364 | -0.606 | 0.126 |
|  | Std. Error | 0.445 | 0.472 | 0.521 | 0.817 | 0.439 | 0.438 | 0.511 | 0.649 | 0.512 | 0.650 | 0.438 | 0.511 | 0.649 |
|  | t value | 15.852 | 4.534 | -3.440 | -1.613 | 0.962 | 0.941 | -0.838 | 1.069 | -0.056 | 0.727 | 0.831 | -1.186 | 0.195 |
|  | Pr(>\|t\|) | **< 0.001** | **< 0.001** | **0.001** | 0.109 | 0.338 | 0.349 | 0.404 | 0.288 | 0.955 | 0.469 | 0.408 | 0.238 | 0.846 |
| EPT taxa | Estimate | 4.760 | 1.603 | -0.948 | -0.658 | -0.061 | -0.160 | -0.155 | 0.481 | 0.214 | 0.052 | 0.118 | -0.422 | -0.385 |
|  | Std. Error | 0.277 | 0.309 | 0.329 | 0.527 | 0.272 | 0.272 | 0.321 | 0.415 | 0.321 | 0.415 | 0.272 | 0.321 | 0.415 |
|  | z value | 17.153 | 5.192 | -2.883 | -1.248 | -0.224 | -0.586 | -0.483 | 1.161 | 0.665 | 0.125 | 0.433 | -1.313 | -0.930 |
|  | Pr(>\|z\|) | **< 0.001** | **< 0.001** | **0.005** | 0.215 | 0.823 | 0.559 | 0.630 | 0.248 | 0.507 | 0.901 | 0.666 | 0.192 | 0.355 |

When analysing the drift behaviour of individual taxa, the microcrustaceans and chironomid larvae showed responses similar to those observed in the drift behaviour of all invertebrates as expected due to their dominance. Predation exhibited a three-way interaction with salinity and temperature (glm, chironomid larvae: df 112, t-value -3.183, Pr(>|t|) 0.002; microcrustacean: df 112, t-value -3.297, Pr(>|t|) < 0.001; Tab. 1). Both groups showed a significant increase of drift behaviour when exposed to predation solely under controlled conditions without the two stressors (emmeans, chironomid larvae: indirect predation: estimate df Inf, z-ratio -2.729, p-value 0.010; direct predation: df Inf, z-ratio -3.080, p-value 0.006; microcrustacean: indirect predation: df Inf, z-ratio -2.368, p-value 0.027; direct predation: df Inf, z-ratio -3.139, p-value 0.005; Supplementary tab. 1), or when both stressors influenced the effects of direct predator (emmeans, direct predation: chironomid larvae: df Inf, z-ratio - 3.177, p-value 0.005; microcrustacean: df Inf, z-ratio -3.553, p-value 0.001; Supplementary tab. 1). Gammarids and EPT taxa were only influenced by direct predation (glm, gammarids: df 112, t-value -3.440, Pr(>|t|) < 0.001; EPT taxa: df 112, t-value -2.883, Pr(>|t|) 0.005; Supplementary tab. 1). The stressors had no direct effect on drift behaviour, except that increased salinity caused more chironomid larvae to drift (emmeans: df Inf, z-ratio -2.092, p-value 0.036; Supplementary tab. 1).

### 3.2 Invertebrates in the leaf litter

The total number of invertebrates, collected from mesh bags filled with leaf litter, remained unchanged by predator exposure, salinity, temperature or their combination (Tab. 2).

**Tab. 2:**
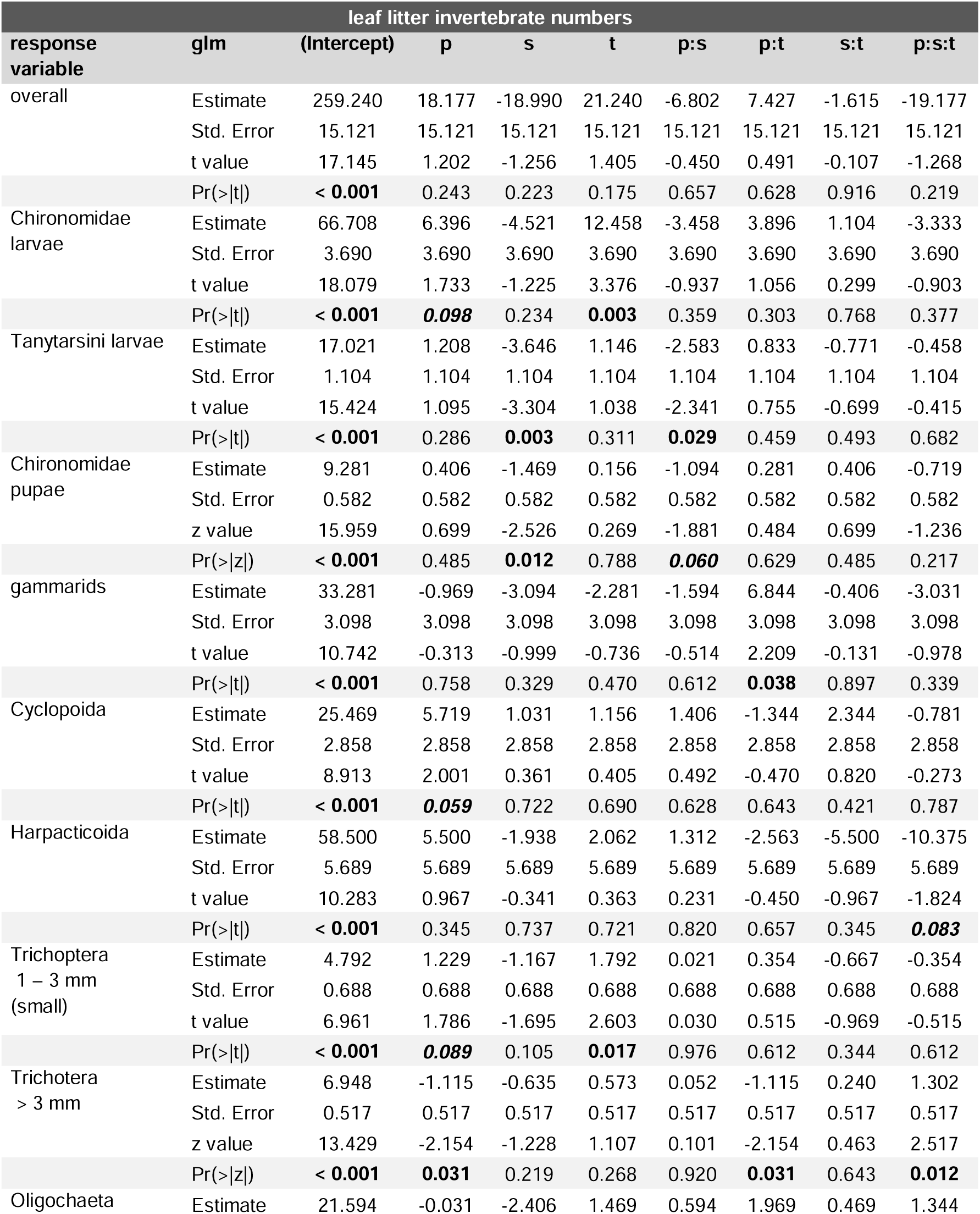
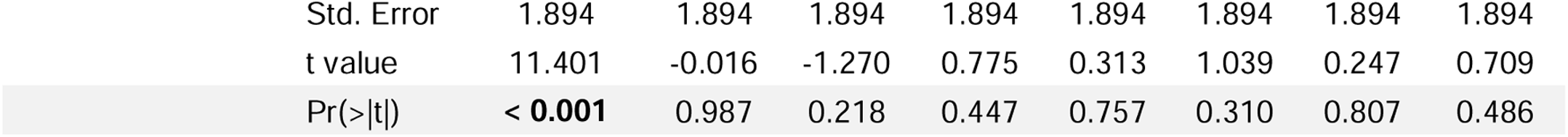
Summary of GLM (generalised linear model) results of invertebrate numbers found in the leaf litter. Given are the effects and interactions of the factors predator exposure (p), salinity (s), and temperature (t). Predator-specific cues were introduced continuously into the system for indirect predator exposure. On the fifth day, predator exposure was augmented by introducing bullheads into the mesocosms. Bold and italic font denotes tendential (< 0.05) and bold font denotes significant (< 0.1) p-values.

| leaf litter invertebrate numbers |  |  |  |  |  |  |  |  |  |
| --- | --- | --- | --- | --- | --- | --- | --- | --- | --- |
| response variable | glm | (Intercept) | p | s | t | p:s | p:t | s:t | p:s:t |
| overall | Estimate | 259.240 | 18.177 | -18.990 | 21.240 | -6.802 | 7.427 | -1.615 | -19.177 |
|  | Std. Error | 15.121 | 15.121 | 15.121 | 15.121 | 15.121 | 15.121 | 15.121 | 15.121 |
|  | t value | 17.145 | 1.202 | -1.256 | 1.405 | -0.450 | 0.491 | -0.107 | -1.268 |
|  | Pr(>\|t\|) | **< 0.001** | 0.243 | 0.223 | 0.175 | 0.657 | 0.628 | 0.916 | 0.219 |
| Chironomidae larvae | Estimate | 66.708 | 6.396 | -4.521 | 12.458 | -3.458 | 3.896 | 1.104 | -3.333 |
|  | Std. Error | 3.690 | 3.690 | 3.690 | 3.690 | 3.690 | 3.690 | 3.690 | 3.690 |
|  | t value | 18.079 | 1.733 | -1.225 | 3.376 | -0.937 | 1.056 | 0.299 | -0.903 |
|  | Pr(>\|t\|) | **< 0.001** | **0.098** | 0.234 | **0.003** | 0.359 | 0.303 | 0.768 | 0.377 |
| Tanytarsini larvae | Estimate | 17.021 | 1.208 | -3.646 | 1.146 | -2.583 | 0.833 | -0.771 | -0.458 |
|  | Std. Error | 1.104 | 1.104 | 1.104 | 1.104 | 1.104 | 1.104 | 1.104 | 1.104 |
|  | t value | 15.424 | 1.095 | -3.304 | 1.038 | -2.341 | 0.755 | -0.699 | -0.415 |
|  | Pr(>\|t\|) | **< 0.001** | 0.286 | **0.003** | 0.311 | **0.029** | 0.459 | 0.493 | 0.682 |
| Chironomidae pupae | Estimate | 9.281 | 0.406 | -1.469 | 0.156 | -1.094 | 0.281 | 0.406 | -0.719 |
|  | Std. Error | 0.582 | 0.582 | 0.582 | 0.582 | 0.582 | 0.582 | 0.582 | 0.582 |
|  | z value | 15.959 | 0.699 | -2.526 | 0.269 | -1.881 | 0.484 | 0.699 | -1.236 |
|  | Pr(>\|z\|) | **< 0.001** | 0.485 | **0.012** | 0.788 | **0.060** | 0.629 | 0.485 | 0.217 |
| gammarids | Estimate | 33.281 | -0.969 | -3.094 | -2.281 | -1.594 | 6.844 | -0.406 | -3.031 |
|  | Std. Error | 3.098 | 3.098 | 3.098 | 3.098 | 3.098 | 3.098 | 3.098 | 3.098 |
|  | t value | 10.742 | -0.313 | -0.999 | -0.736 | -0.514 | 2.209 | -0.131 | -0.978 |
|  | Pr(>\|t\|) | **< 0.001** | 0.758 | 0.329 | 0.470 | 0.612 | **0.038** | 0.897 | 0.339 |
| Cyclopoida | Estimate | 25.469 | 5.719 | 1.031 | 1.156 | 1.406 | -1.344 | 2.344 | -0.781 |
|  | Std. Error | 2.858 | 2.858 | 2.858 | 2.858 | 2.858 | 2.858 | 2.858 | 2.858 |
|  | t value | 8.913 | 2.001 | 0.361 | 0.405 | 0.492 | -0.470 | 0.820 | -0.273 |
|  | Pr(>\|t\|) | **< 0.001** | **0.059** | 0.722 | 0.690 | 0.628 | 0.643 | 0.421 | 0.787 |
| Harpacticoida | Estimate | 58.500 | 5.500 | -1.938 | 2.062 | 1.312 | -2.563 | -5.500 | -10.375 |
|  | Std. Error | 5.689 | 5.689 | 5.689 | 5.689 | 5.689 | 5.689 | 5.689 | 5.689 |
|  | t value | 10.283 | 0.967 | -0.341 | 0.363 | 0.231 | -0.450 | -0.967 | -1.824 |
|  | Pr(>\|t\|) | **< 0.001** | 0.345 | 0.737 | 0.721 | 0.820 | 0.657 | 0.345 | **0.083** |
| Trichoptera<br>1 – 3 mm<br>(small) | Estimate | 4.792 | 1.229 | -1.167 | 1.792 | 0.021 | 0.354 | -0.667 | -0.354 |
|  | Std. Error | 0.688 | 0.688 | 0.688 | 0.688 | 0.688 | 0.688 | 0.688 | 0.688 |
|  | t value | 6.961 | 1.786 | -1.695 | 2.603 | 0.030 | 0.515 | -0.969 | -0.515 |
|  | Pr(>\|t\|) | **< 0.001** | **0.089** | 0.105 | **0.017** | 0.976 | 0.612 | 0.344 | 0.612 |
| Trichoptera<br>> 3 mm | Estimate | 6.948 | -1.115 | -0.635 | 0.573 | 0.052 | -1.115 | 0.240 | 1.302 |
|  | Std. Error | 0.517 | 0.517 | 0.517 | 0.517 | 0.517 | 0.517 | 0.517 | 0.517 |
|  | z value | 13.429 | -2.154 | -1.228 | 1.107 | 0.101 | -2.154 | 0.463 | 2.517 |
|  | Pr(>\|z\|) | **< 0.001** | **0.031** | 0.219 | 0.268 | 0.920 | **0.031** | 0.643 | **0.012** |
| Oligochaeta | Estimate | 21.594 | -0.031 | -2.406 | 1.469 | 0.594 | 1.969 | 0.469 | 1.344 |

|  |  |  |  |  |  |  |  |  |
| --- | --- | --- | --- | --- | --- | --- | --- | --- |
| Std. Error | 1.894 | 1.894 | 1.894 | 1.894 | 1.894 | 1.894 | 1.894 | 1.894 |
| t value | 11.401 | -0.016 | -1.270 | 0.775 | 0.313 | 1.039 | 0.247 | 0.709 |
| Pr(>\|t\|) | **< 0.001** | 0.987 | 0.218 | 0.447 | 0.757 | 0.310 | 0.807 | 0.486 |

We found a trend of predation reducing the abundance of some taxa (Cyclopoida: df 21, t-value 2.001, Pr(>|z|) 0.059; Chironomidae larvae: df 21, t-value 1.733, Pr(>|z|) 0.098; small Trichoptera: df 21, t-value 1.786, Pr(>|z|) 0.089; Tab. 2). Salinity significantly increased the number of Tanytarsini larvae and Chironomidae pupae (Tanytarsini larvae, emmeans: df Inf, z-value -3.974, p-value < 0.001, Supplementary tab. 2; Chironomidae pupae, glm: t-value -2.526, Pr(>|z|) 0.012; Tab. 2). Increased temperature decreased the number of Chironomidae larvae (glm: df 21, t-value 3.376, Pr(>|z|) 0.003) and small Trichoptera (glm: t-value 2.603, Pr(>|z|) 0.017; Tab. 2). Oligochaeta remained unaffected across all treatments.

Predation and increased salinity or temperature affected distinct individual taxa. Specifically, the effect of increased salinity on the number of Tanytarsini larvae was diminished when exposed to predation (glm: df 21, t-value -2.341, Pr(>|z|) 0.029; Tab. 2). Increased individual numbers of Tanytarsini larvae were observed only in response to elevated salinity levels (emmeans: df Inf, z-value -3.974, p-value < 0.001; Supplementary tab. 2). Similarly, in the Chironomidae pupae we found that predator exposure and salinity tended to interact antagonistic with individual numbers only increasing when exposed to increased salinity without predation (glm: df 21, t-value -1.881, Pr(>|z|) 0.060; Tab. 2). In contrast, the significant antagonistic interaction between predator exposure and temperature increased the individual numbers of gammarids (glm: df 21, t-value 2.209, Pr(>|z|) 0.038; Tab. 2).

Predation, salinity, and temperature had a significant three-way interaction effect on Trichoptera larger than 3 mm (glm: df 21, z-value 2.517, Pr(>|z|) 0.012; Tab. 2). Increased individual numbers in the leaf litter due to predator exposure were observed only under elevated salinity conditions (emmeans salinity vs. predation salinity: df Inf, z-ratio -2.672, p-value 0.008; Supplementary tab. 2). This increase due to salinity and predation was reduced by increased temperature (emmeans predation salinity vs. predation salinity temperature: df Inf, z-ratio 2.043, p-value 0.041; Tab. 2). We found a tendentially three-way interaction increasing the number of Harpacticoida (glm: df 21, t-value -1.824, Pr(>|z|) 0.083; Tab. 2).

### 3.3 Invertebrates in the community

The invertebrates found in leaf litter and channel substrate were considered to be part of the remaining community as they stayed in the system. They were combined to analyse how salinization and warming interacted with the effects of predation on the entire community. Alone predation (glm: df 21, t-value 2.076, Pr(>|z|) 0.050) and increased temperature (glm: df 21, t-value 2.419, Pr(>|z|) 0.025) significantly reduced the total number of individuals compared to ambient conditions (Fig. 6, Tab. 3). We did not observe any interaction between predation, increased salinity and increased temperature.

**Fig. 6:**
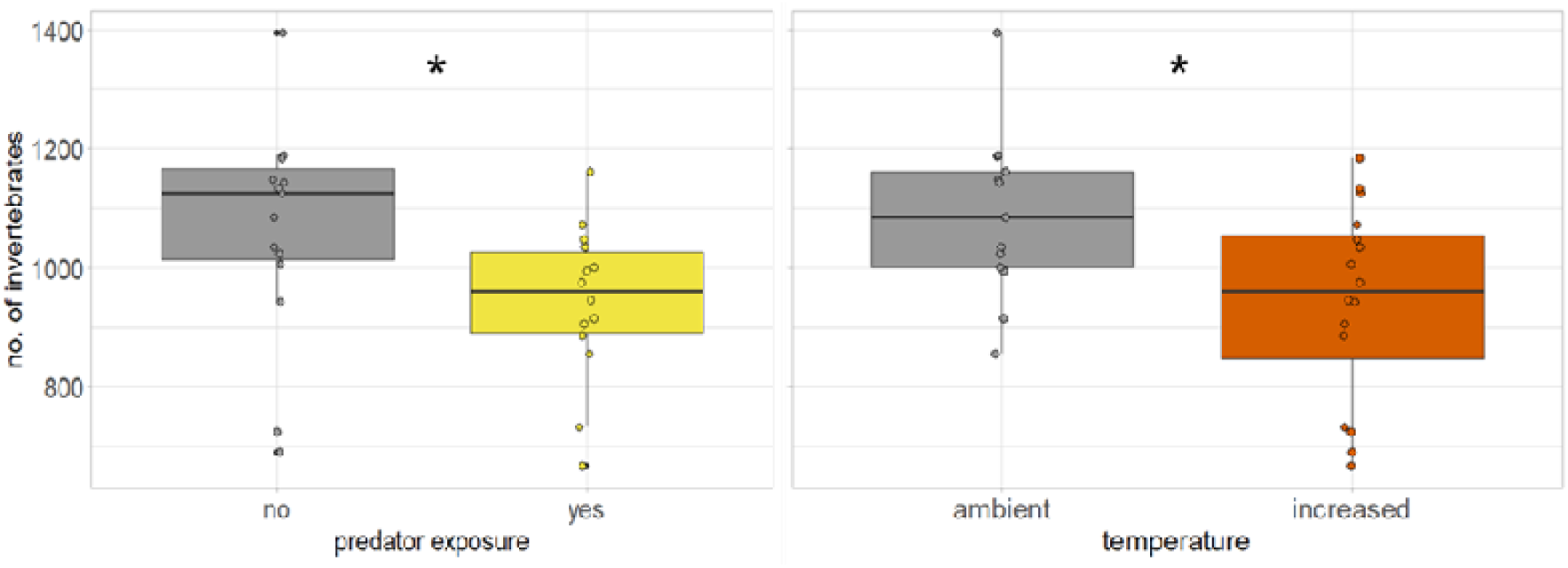
Number of invertebrates staying in the system after exposure to predation pressure or elevated temperature. The communities were combined from invertebrates found in leaf litter and substrate. Predator exposure was caused by the fish species *Gasterosteus aculeatus* and *Cottus rhenanus*. Both predator exposure (Estimate 58.969, Std. Error 28.410, t-value 2.076, Pr(>|t|) 0.050) and temperature (Estimate 68.719, Std. Error 28.410, t value 2.419, Pr(>|t|) 0.025) led to a reduction of the invertebrate communities numbers. Data was fitted with a generalised linear model (glm).

**Tab. 3:**
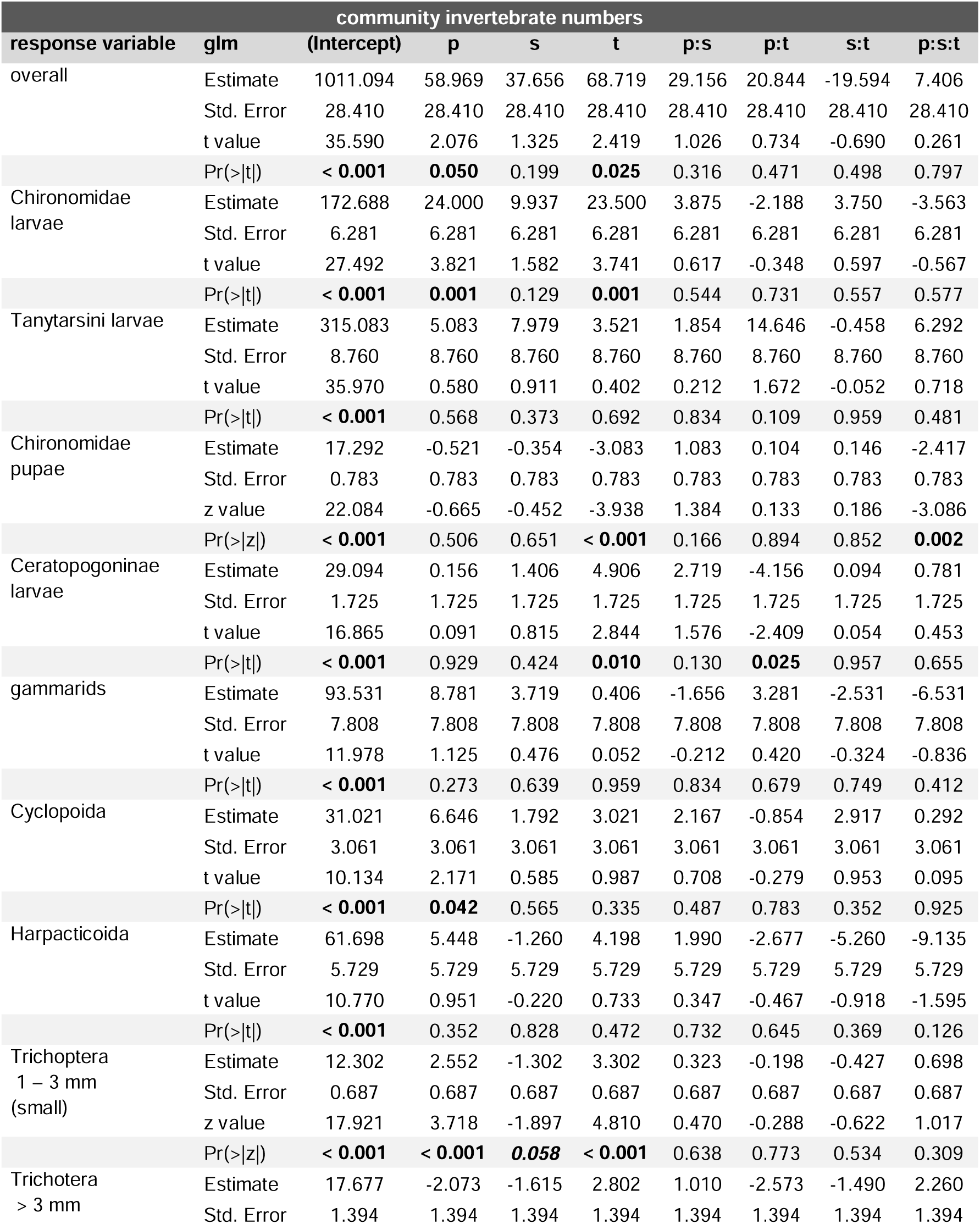
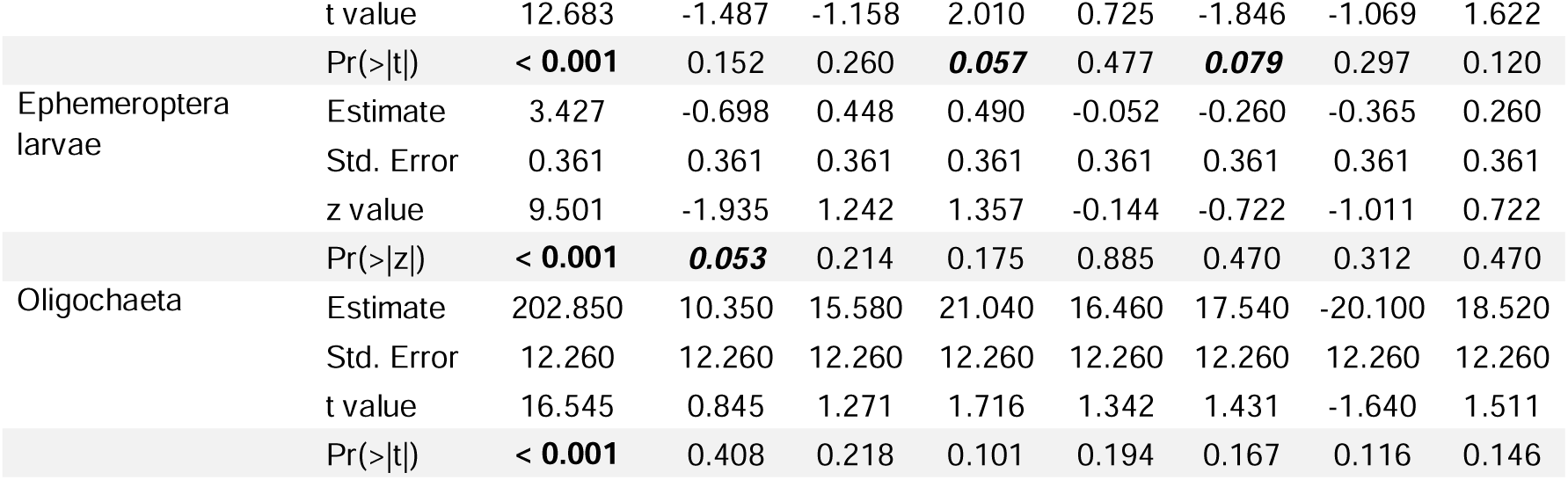
Summary of GLM (generalized linear model) results of invertebrate numbers of the community. Given are the effects and interactions of the factors predator exposure (p), salinity (s), and temperature (t). Predator-specific cues were introduced continuously into the system for indirect predator exposure. On the fifth day, predator exposure was augmented by introducing bullheads into the mesocosms. Bold and italic font denotes tendential (< 0.05) and bold font denotes significant (< 0.1) p-values.

Investigating individual taxa, gammarids, Tanytarsini larvae, Harpacticoida and Oligochaeta showed no response to any treatment (Tab. 3). Predator exposure affected the Cyclopoida (glm: df 21, t-value 2.171, Pr(>|z|) 0.042), small Trichoptera (glm: df 21, t-value 3.718, Pr(>|z|) < 0.001) and Chironomidae larvae (glm: df 21, t-value 3.821, Pr(>|z|) 0.001), resulting in reduced individual numbers (Tab. 3). There was a trend towards higher individual numbers of Ephemeroptera larvae upon predation (glm: df 21, t-value -1.935, Pr(>|z|) 0.053; Tab. 3). Populations of Chironomidae larvae and small Trichoptera were reduced under elevated temperature (glm Chironomidae larvae: df 21, t-value 3.741, Pr(>|z|) 0.001; small Trichoptera: df 21, t-value 4.810, Pr(>|z|) <0.001; Tab. 3). Salinity tendentially increased the number of small Trichoptera individuals (glm: df 21, t-value -1.897, Pr(>|z|) 0.058). No influence of increased salinity was observed on any other taxa (Tab. 3).

The individual numbers of Ceratopogonidae larvae were significantly reduced by the synergistic interaction of increased temperature and predation (glm: df 21, t-value -2.409, Pr(>|z|) 0.025; Tab. 3). Predation led to a decrease in the number of Ceratopogonidae larvae when temperatures were elevated (emmeans, predation vs. predation+temperature: df Inf, z-ratio 3.529 p-value < 0.001; Supplementary tab. 3). Trichoptera over 3 mm in size were also tendentially reduced by the synergistic interaction of increased temperature and predator exposure (glm: df 21, z-value -1.846, Pr(>|z|) 0.079; Tab. 3). Temperature significantly reduced the individual numbers of Trichoptera over 3 mm (emmeans: df Inf, z-value -2.042, p-value 0.041; Supplementary tab. 3). There was no interaction between predation and salinity on individual taxa numbers (Tab. 3).

A significant three-way interaction between predation, salinity, and temperature affected the number of Chironomidae pupae (glm: df 21, t-value -3.086, Pr(>|z|) 0.002; Tab. 3) with the temperature stressor influencing the interaction between fish and salinity, resulting in an increase in pupae numbers (emmeans, predation+salinity vs predation+salinity+temperature: df Inf, z-ratio -3.190, p-value 0.001; Supplementary tab. 3). Elevated temperature increased pupae numbers (emmeans, control vs. temperature: df Inf, z-ratio -3.550, p-value < 0.001; Supplementary tab. 3).

## 4 Discussion

### 4.1 Predator-induced behaviour

We here demonstrated that fish exposure in riverine ecosystems triggered **predator-induced drift behaviour** in invertebrate prey. This behaviour occurred within the first 24 hours of each predation stimulus (i.e., indirect and direct predator exposure). The observed response on the first day likely relied on chemical cues, as the prey taxa were not in direct contact with the predators (Brönmark and Hansson, 2012, 2000; Tollrian and Harvell, 1999).

Chironomids and microcrustaceans detected and responded to both indirect and direct predation, whereas gammarids and EPT taxa only exhibited increased drift behaviour in response to direct predation by bullheads. Previous studies showed that predation typically results in reduced activity across most taxa (Åbjörnsson et al., 1997; Ahlgren et al., 2011; Gall and Brodie, 2009; Huryn and Chivers, 1999; Kasumyan, 2022; Schäffer et al., 2013; Williams and Moore, 1982; Wisenden et al., 1997), although some, particularly Baetidae, also exhibited increased drift and movement (Culp et al., 1991; Lancaster, 1990; Poff et al., 1991). Similarly, studies found microcrustaceans either decreasing their activity (Bjærke et al., 2016; Heuschele et al., 2020) or seeking refuge through vertical and horizontal migration (Jack et al., 2006; Lauridsen and Lodge, 1996; Pasternak et al., 2006; Weiss and Tollrian, 2018; Zaret and Suffern, 1976). However, many of these studies focused on only a few select taxa with limited numbers of individuals, potentially obscuring density and biotic interaction effects (Peacor and Werner, 1997; Tollrian et al., 2015). Additionally, most of the studies were primarily conducted in systems without water currents needed to study drifting. The differences in experimental setups may explain the discrepancies between our and previous observations. In comparison, Dahl (1998b) also documented increased drift rates for *Gammarus*, *Ephemerella*, and *Baetis* when studying the effects of a fish predator on an invertebrate community.

While all tested invertebrate taxa in our study showed increased drift behaviour, variations in the timing of initiation could reflect differences in their perception of predation risk. These differences may be influenced by factors such as the predator species and diet, predator and prey density, the prey sensitivity and the level of predation cue concentration (Brown et al., 2006; Dahl, 1998b; Huryn and Chivers, 1999; Peckarsky and McIntosh, 1998; Stabell et al., 2003; Tollrian et al., 2015). Gammarids and EPT taxa, which responded exclusively to direct predation, may as such prioritize cryptic behaviour over immediate flight via drifting when detecting only cues, particularly when the perceived threat level is not sufficiently high. It is only under the high threat level of direct predation, where cue concentration is further increased and taxa can also visually detect and directly encounter the predator, that their strategies would then be adjusted, triggering the drift response. The threat sensitivity hypothesis states that prey species adjust their anti-predator behaviour in response to varying levels of perceived threat to minimise costs (Helfman, 1989; Sih, 1980). Indeed, drift behaviour is a strategic adaptation that tends to evolve in environments with temporary or spatially heterogeneous predator abundance (Tollrian and Harvell, 1999). As such, this flexibility in behaviour observed in Gammarids and EPT taxa may indicate their ability to adjust their response to varying levels of risk, minimizing energetic costs.

We did not observe changes in invertebrate numbers in the leaf litter. The presence of invertebrates in the leaf litter bags could indicate either foraging activity or hiding behaviour. However, under predation, we would have interpreted increased invertebrate numbers as hiding behaviour, as predation typically increases hiding and reduces foraging activity. (Alvarez et al., 2014; Kasumyan, 2022; Lauridsen and Lodge, 1996). As such based on the similar overall number of individuals in the leaf litter in the control and predator conditions **predator-induced hiding behaviour** in the leaf litter did not take place. Only a few taxa tendentially displayed hiding behaviour in response to predation with slightly higher numbers of individuals in the fish treatment. This may depend on the high predation risk (e.g., 1 fish per mesocosm with 3.5 L volume) that we applied. It is possible that the threat level surpassed a certain threshold, prompting invertebrates to opt for a direct avoidance strategy through fleeing the system by drift (Helfman, 1989). Alternatively, the leaf litter bags may not have been attractive refuges for certain taxa, leading the invertebrates to seek other shelters such as sediment and interstitial spaces.

Otherwise, it can also be discussed that by the end of the 14 days with continuous predation risk, the predator-induced hiding behaviour had already ceased, and prey species continued foraging outside the leaf litter. This has, e.g., been observed in mayfly larvae that exhibited only increased short-term drifting behaviour in response to predators (Culp et al., 1991). In fact, many prey animals balance the costs of predator-induced behaviour against other fitness-related activities such as foraging and mating (Bjærke et al., 2016; Lima and Dill, 1990; Sih, 1980). This trade-off often limits the duration of predator-induced behaviour.

### 4.2 Stressor-induced behaviour

Salinization and warming are known to affect invertebrate behaviour, as observed in the higher drift rates of sensitive invertebrate taxa (Beermann et al., 2018; Wood and Dykes, 2002). In our study, only the chironomid larvae exhibited increased drift due to elevated salinity, despite the general ability of Chironomidae to thrive across a wide range of salinity concentrations (Golovatyuk et al., 2022; Zinchenko and Golovatyuk, 2013). The EPT taxa, which are often more sensitive (Zinchenko and Golovatyuk, 2013) showed no drift reaction to elevated salt and temperature conditions. The lack of response from sensitive taxa may stem from the ‘stressor legacy’ (Jackson et al., 2021; Padisak, 1992) of historic coal mining in the tested stream and its community. Long-term exposure of this community to high salt levels could explain its differing response to the current stressors (Lee and Petersen, 2003; Madge Pimentel et al., 2024a; Schröder et al., 2015). For example, increased salinity may have led to shifts in community composition, with sensitive taxa either adapting or departing, leaving behind only resistant taxa. This could have bolstered the community’s survivability and reduced their drift response in the face of heightened salinity levels. We cannot entirely discount the possibility that salinity still affects the invertebrates in other ways (such as growth rate or emergence time), which were not measured in our study. That chironomids reacted despite the possibility of adaptation due to stressor legacy could be attributed to, e.g., their multivoltine life cycles, allowing for rapid generational turnover (Drake, 1982; Pinder, 1986). Once the stream was restored, this adaptation could therefore fade rapidly, resulting in an observable drift response under elevated salinity conditions. The presence of a stressor legacy and the potential for rapid decline in adaptation require further investigation.

Similarly, temperature increase alone did not affect the drift behaviour of the invertebrates. This may not necessarily indicate adaptation, but rather the magnitude of temperature change. With only a 3.4-degree difference outside of a heatwave, the mesocosm temperatures ranged from a minimum of 5.2°C to a maximum of 16.2°C with a mean of 10.1 °C ± 2.4°C. The temperature change may simply not have exceeded the thermal tolerance range of the taxa (Dallas and Ketley, 2011; Quinn et al., 1994).

Invertebrate numbers in the leaf litter were also largely unaffected by the stressors, with only some taxa showing changes. Under increased salinity conditions, Tanytarsini larvae and Chironomidae pupae were more commonly found in the leaf litter, contradicting our expectation of decreased numbers due to negative effects. It is unlikely that this increased presence indicates traditional hiding behaviour, as remaining in the leaf litter does not offer refuge from the elevated salinity levels. Considering that Tanytarsini larvae are known for their resilience to elevated salinity levels (Dimitriadis and Cranston, 2007; Zinchenko and Golovatyuk, 2013) and chironomids lack osmotic regulation in the pupal stage (Berezina, 2003), thus likely delaying pupation in response to salinity stress, it is unlikely that they sought refuge in the leaf litter. This implies that the salinity levels used in our study may not have had a significant impact on the behaviour and could even have been beneficial.

Conversely, Chironomidae larvae and smaller Trichoptera were less frequently found within the leaf litter under increased temperature conditions. This indicates a reduction in their time spent in the leaf litter due to the heightened metabolic demands. With metabolism being enhanced by increased temperatures, there is also a greater demand for energy intake (Heiman and Knight, 1975; Suarez, 2012). They could have left in search of resources that are more easily assimilated, to find higher quality food, as a thermoregulatory behaviour to adjust their body temperature, or to seek thermal refugia by migrating to cooler water areas (Berman and Quinn, 1991; Gordon et al., 2018). Otherwise, the oxygen availability could have been too low due to temperature and decomposition processes, forcing them to flee. This is unlikely, though, as most taxa tolerate all but the very lowest DO levels (Connolly et al., 2004).

### 4.3 Disruption of predator-induced drift behaviour by stressors

In our study, the effect of predation on drift was hampered when invertebrates were exposed to increased salinity or increased temperature. These stressors not only affected the overall number of invertebrates displaying predator-induced drift behaviour but also individual taxa, i.e., chironomid larvae and microcrustaceans. We observed this effect both in response to indirect and direct predation. Other studies have reported a comparable reduction of response to predator cues under stressor conditions due to their effect on cue concentration or the prey’s sensing ability (Trotter et al., 2019; Van Donk et al., 2015; Weiss et al., 2018).

We could not fully confirm that the combination of increased salinity and temperature also suppresses predator-induced drift behaviour. Although the stressors did decrease the response to indirect predation, direct predation still resulted in an increased number of drifting invertebrates. The reasons for this effect are unclear as both single stressors fully suppressed the predator-induced behaviour. It is possible that direct predation induces more stress due to higher cue concentration, visual cues and the imminent danger. In such cases, the interaction of direct predation risk and the stressors might have exceeded the invertebrates’ tolerance level, leading to emigration (Howard and Webster, 2009; Schäfer and Piggott, 2018).

As we did not observe predator-induced hiding behaviour, there was no response to be altered by the stressors.

### 4.4 Altered predator-induced drift behaviour despite salinity adaptation

The limited response of invertebrates to increased salinity contrasts sharply with its significant influence on predator-induced behaviour, revealing a noteworthy paradox: while the drift behaviour of prey organisms appeared unaffected by salinity, its presence still shapes predator-prey interactions. Sections of the Boye River were contaminated with wastewater until 2017, with maximum chloride levels reaching up to 15,500 mg/L in 1996 (www.elwasweb.nrw.de, last accessed April 19, 2024). These decades of exposure to poor conditions would have forced sensitive taxa to adapt either through physiological or behavioural adjustments or be replaced by more tolerant species. The salinity adaptation of the invertebrates likely mitigated the impact of the stressors. However, the sudden experimental increase in salinity may still cause physiological stress, necessitating acclimation to the new condition. Even in euryhaline species, this process involves trade-offs (Tietze and Gerald, 2016). Increased salinity leads to the reallocation of energy to essential physiological processes, such as the expression of detoxification enzymes and ion transporter genes, which regulate the membrane permeability of sodium, potassium, and chloride (Brasseur et al., 2022). These adjustments may enable invertebrates to better withstand salinity stress, but they might compromise other life relevant aspects such as behaviour. Physiological trade-offs could constrain invertebrates’ ability to maintain optimal behavioural responses under salinity stress.

Moreover, as acclimation to saline conditions likely prioritises cellular equilibrium and survival, higher salinity may continue to adversely affect the sensory system for predator recognition and evasion (Ross and Behringer, 2019). Consequently, despite surviving in saline environments, invertebrates may remain susceptible due to inadequate modulations of adaptive behaviours.

Even without considering its direct effects on invertebrates in freshwater, salinity can influence pH, conductivity, solubility, ion exchange and nutrient mobilization (Haq et al., 2018; Löfgren, 2001). These factors could potentially disrupt molecular interactions, altering compound properties, and affecting the function or concentration of chemical cues in the water. Such alterations could potentially hinder the taxa’s ability to sense predators despite their adaptations.

### 4.5 Effects of predation and stressors on community

On the community level, invertebrate numbers were only affected by predation and increased temperature. Predation reduced individual numbers in the community partly due to the increased predator-induced drift behaviour through which taxa fled from the environment. This is consistent with previous mesocosm experiments showing an inverse relationship between drift behaviour and final abundance (Beermann et al., 2018; Magbanua et al., 2016; Piggott et al., 2015). Additionally, direct predation might have also contributed to the reduction in invertebrate numbers as a result of prey consumption or higher mortality rates due to stress.

In response to increased temperature, overall invertebrate numbers (e.g., Chironomidae and Trichoptera larvae) decreased, which was furthermore in line with an increase in the Chironomidae pupae counts. This suggests that the temperature increase of 3.4 °C accelerated insect development. A similar pattern was observed by Hogg and Williams (1996), where Chironomidae densities decreased and an earlier onset of adult insect emergence occurred. Considering the mild temperatures observed during our study in March, it is unlikely that the temperature increase exceeded the tolerance range at any point of the invertebrates (Dallas and Ross-Gillespie, 2015; Quinn et al., 1994), thus minimising the likelihood of a negative impact. In contrast, salinization did not affect the number of invertebrates in the community, possibly due to the “stressor legacy” of historic coal mining. Having likely already adapted to higher degrees of salinity, the chosen salinity level, based on the known µS/cm maxima of the Boye catchment, may not have exceeded their elevated tolerance level.

We had anticipated an even stronger decrease in individuals under direct predation when the stressors influenced predator-induced responses. Yet, no interaction between predation and the stressors was observed. This suggests that invertebrates were not significantly more consumed. The foremost reason for the minimal direct predation effect may lie in the short duration of the bullheads’ presence in the mesocosm, as several left within nine days, likely consuming less than expected. We anticipate that predators could experience greater hunting success when prolonged exposure to stressors leads to altered behavioural responses in invertebrates, thereby impacting the invertebrate community.

## 5 Conclusion

In conclusion, we were able to use the *ExStream* system to successfully study predator-induced behaviour. While implementing indirect predation functioned well, direct predation led to methodological inconsistencies in our experimental setup.

We demonstrated that predation induces increased drift behaviour in a range of invertebrate taxa, impacting the overall community. Anthropogenic stressors, including salinity, temperature, and their combination, partially inhibited predator-induced drift behaviour. Despite the potential adaptation to higher stressor levels from prolonged exposure in the Boye, invertebrate behaviour was still altered. In contrast, hiding behaviour remained unchanged, possibly due to varying defence strategies or because any predator-induced behaviour had subsided by the time of sampling. Additionally, the community also stayed largely unaffected, with only predation and temperature influencing the overall invertebrate numbers and no interaction observed. The results of both the hiding behaviour and the community numbers were likely influenced by the small sample size and short duration of the study, with treatment differences possibly being too subtle for our analysis to detect differences.

These findings suggest that invertebrates in riverine ecosystems subjected to stressors may exhibit a reduced ability to respond appropriately to predators, as evidenced by the changes in drift behaviour. While this did not immediately affect our community within the two-week timeframe, it may have long-term consequences not observed in our study. In natural environments, we anticipate that such altered responses combined with direct encounters with fish predators, increase the fish’s hunting success rates.

## 6 Acknowledgements

A sincere thank you is owed to Dr Christian Edler, Birgit Daniel (Bezirksregierung Düsseldorf, Dezernat 51) and Bernd Stemmer (Bezirksregierung Arnsberg, Fischereidezernat) for their invaluable assistance in catching the fish. We also thank Christoph D. Matthaei for his valuable advice. Special thanks go to the many people who actively supported us during the mesocosm experiment, including Alexander Rogalla, Theodore Domke, Antonia Domnik, Alexandra Hollstein, Johanna Knupfer, and Melina Rüngeler. We also wish to acknowledge and extend our sincere gratitude to Nicolai Bissantz and Willem Kaijser for their assistance and expertise in addressing our statistical inquiries. This project is part of the Collaborative Research Centre 1439 RESIST (Multilevel Response to Stressor Increase and Decrease in Stream Ecosystems; www.sfb-resist.de) funded by the Deutsche Forschungsgemeinschaft (DFG, German Research Foundation; CRC 1439/1, project number: 426547801).

## Supplementary files

Supplementary fig. 1: Conductivity fluctuations of the stressor mesocosms during the 14-day stressor phase of the *ExStream* system.

Supplementary fig. 2: Temperature fluctuations during the stressor phase of 14 days.

Supplementary tab. 1: Summary of post hoc comparisons among groups results (emmeans).

Supplementary tab. 2: Summary of post hoc comparisons among groups results (emmeans).

Supplementary tab. 3: Summary of post hoc comparisons among groups results (emmeans).

## Statement of authorship

The project was planned by RT. The Initial planning for the implementation of the mesocosm experiment was done by FL, AJB and RT. They also organized the necessary administrative communication. Resources were organised by IMP. AMV, IMP, and PMR implemented and improved the system. AMV investigated predator-specific behaviour. AMV, IMP, AJB, MH and LCW contributed to the statistical analysis of the data. AMV and TO processed the samples and analysed the data and AMV wrote the initial draft of the manuscript. All authors contributed to the draft and approved the final version of the manuscript.

## References

Åbjörnsson, K., Wagner, B.M.A., Axelsson, A., Bjerselius, R., Olsén, K.H., 1997. Responses of Acilius sulcatus (Coleoptera: Dytiscidae) to chemical cues from perch (Perca fluviatilis). Oecologia 111, 166–171. 10.1007/s004420050221

Ahlgren, J., Åbjörnsson, K., Brönmark, C., 2011. The influence of predator regime on the behaviour and mortality of a freshwater amphipod, Gammarus pulex. Hydrobiologia 671, 39–49. 10.1007/s10750-011-0702-8

Alvarez, M., Landeira-Dabarca, A., Peckarsky, B., 2014. Origin and specificity of predatory fish cues detected by Baetis larvae (Ephemeroptera; Insecta). Anim. Behav. 96, 141–149. 10.1016/j.anbehav.2014.07.017

Bauernfeind, E., Humpesch, U., 2001. Die Eintagsfliegen Zentraleuropas (Insecta: Ephemeroptera): Bestimmung und Ökologie. Verlag des Naturhistorischen Museums Wien.

Beermann, A.J., Elbrecht, V., Karnatz, S., Ma, L., Matthaei, C.D., Piggott, J.J., Leese, F., 2018. Multiple-stressor effects on stream macroinvertebrate communities: A mesocosm experiment manipulating salinity, fine sediment and flow velocity. Sci. Total Environ. 610–611, 961–971. 10.1016/j.scitotenv.2017.08.084

Berezina, N.A., 2003. Tolerance of freshwater invertebrates to changes in water salinity. Russ. J. Ecol. 34, 296–301. 10.1023/A:1024597832095

Berman, C.H., Quinn, T.P., 1991. Behavioural thermoregulation and homing by spring chinook salmon, Oncorhynchus tshawytscha (Walbaum), in the Yakima River. J. Fish Biol. 39, 301–312. 10.1111/j.1095-8649.1991.tb04364.x

Bjærke, O., Andersen, T., Bækkedal, K.S., Nordbotten, M., Skau, L.F., Titelman, J., 2016. Paternal energetic investments in copepods. Limnol. Oceanogr. 61, 508–517. 10.1002/lno.10229

Brasseur, M. V., Beermann, A.J., Elbrecht, V., Grabner, D., Peinert-Voss, B., Salis, R., Weiss, M., Mayer, C., Leese, F., 2022. Impacts of multiple anthropogenic stressors on the transcriptional response of Gammarus fossarum in a mesocosm field experiment. BMC Genomics 23, 1–12. 10.1186/s12864-022-09050-1

Brönmark, C., Hansson, L.A., 2012. Chemical Ecology in Aquatic Systems, online edn. Oxford University Press. 10.1093/acprof:osobl/9780199583096.001.0001

Brönmark, C., Hansson, L.A., 2000. Chemical communication in aquatic systems: An introduction. Oikos 88, 103–109. 10.1034/j.1600-0706.2000.880112.x

Brown, G.E., Rive, A.C., Ferrari, M.C.O., Chivers, D.P., 2006. The dynamic nature of antipredator behavior: Prey fish integrate threat-sensitive antipredator responses within background levels of predation risk. Behav. Ecol. Sociobiol. 61, 9–16. 10.1007/s00265-006-0232-y

Buchwalter, D., Scheibener, S., Chou, H., Soucek, D., Elphick, J., 2019. Are sulfate effects in the mayfly Neocloeon triangulifer driven by the cost of ion regulation? Philos. Trans. R. Soc. B Biol. Sci. 374, 2–8. 10.1098/rstb.2018.0013

Cañedo-Argüelles, M., Kefford, B., Schäfer, R., 2019. Salt in freshwaters: Causes, effects and prospects - Introduction to the theme issue. Philos. Trans. R. Soc. B Biol. Sci. 374. 10.1098/rstb.2018.0002

Caro, T., 2014. Antipredator deception in terrestrial vertebrates. Curr. Zool. 60, 16–25. 10.1093/czoolo/60.1.16

Castillo, A.M., Sharpe, D.M.T., Ghalambor, C.K., De León, L.F., 2018. Exploring the effects of salinization on trophic diversity in freshwater ecosystems: a quantitative review. Hydrobiologia 807, 1–17. 10.1007/s10750-017-3403-0

Connolly, N.M., Crossland, M.R., Pearson, R.G., 2004. Effect of low dissolved oxygen on survival, emergence, and drift of tropical stream macroinvertebrates. J. North Am. Benthol. Soc. 23, 251–270. 10.1899/0887-3593(2004)023<0251:EOLDOO>2.0.CO;2

Crowder, L.B., Cooper, W.E., 1982. Habitat Structural Complexity and the Interaction Between Bluegills and Their Prey. Ecol. Soc. Am. 63, 1802–1813.

Culp, J.M., Glozier, N.E., Scrimgeour, G.J., 1991. Reduction of predation risk under the cover of darkness: Avoidance responses of mayfly larvae to a benthic fish. Oecologia 86, 163–169. 10.1007/BF00317527

Dahl, J., 1998a. Effects of a benthivorous and a drift-feeding fish on a benthic stream assemblage. Oecologia 116, 426–432. 10.1007/s004420050606

Dahl, J., 1998b. The impact of vertebrate and invertebrate predators on a stream benthic community. Oecologia 117, 217–226. 10.1007/s004420050651

Dallas, H.F., Ketley, Z.A., 2011. Upper thermal limits of aquatic macroinvertebrates: Comparing critical thermal maxima with 96-LT50 values. J. Therm. Biol. 36, 322–327. 10.1016/j.jtherbio.2011.06.001

Dallas, H.F., Ross-Gillespie, V., 2015. Sublethal effects of temperature on freshwater organisms, with special reference to aquatic insects. Water SA 41, 712–726. 10.4314/wsa.v41i5.15

Dettner, K., 2019. Defenses of Water Insects, in: Del-Claro, K., Guillermo, R. (Eds.), Aquatic Insects. Springer-Nature. 10.1007/978-3-030-16327-3

Dimitriadis, S., Cranston, P.S., 2007. From the mountains to the sea: Assemblage structure and dynamics in Chironomidae (Insecta: Diptera) in the Clyde River estuarine gradient, New South Wales, south-eastern Australia. Aust. J. Entomol. 46, 188–197. 10.1111/j.1440-6055.2007.00592.x

Drake, C.M., 1982. Seasonal dynamics of Chironomidae (Diptera) on the Bulrush Schoenoplectus lacustris in a chalk stream. Freshw. Biol. 12, 225–240. 10.1111/j.1365-2427.1982.tb00618.x

Draper, A.M., Weissburg, M.J., 2019. Impacts of global warming and elevated CO2 on sensory behavior in predator-prey interactions: A review and synthesis. Front. Ecol. Evol. 7. 10.3389/fevo.2019.00072

Eiseler, B., 2010. Taxonomie für die Praxis: Bestimmungshilfen - Makrizoobenthos (1). Landesamt für Natur, Umwelt und Verbraucherschutz Nordrhein-Westfalen, Rechlinghausen.

Eiseler, B., Hess, M., 2015. Taxonomie für die Praxis: Bestimmungshilfen - Makrozoobenthos (2). Landesamt für Natur, Umwelt und Verbraucherschutz Nordrhein-Westfalen, Recklinghausen.

Elbrecht, V., Beermann, A.J., Goessler, G., Neumann, J., Tollrian, R., Wagner, R., Wlecklik, A., Piggott, J.J., Matthaei, C.D., Leese, F., 2016. Multiple-stressor effects on stream invertebrates: A mesocosm experiment manipulating nutrients, fine sediment and flow velocity. Freshw. Biol. 61, 362–375. 10.1111/fwb.12713

Elliott, J.M., Elliott, J.A., 1995. The critical thermal limits for the bullhead, Cottus gobio, from three populations in north-west England. Freshw. Biol. 33, 411–418. 10.1111/j.1365-2427.1995.tb00403.x

Ferrari, M.C.O., Wisenden, B.D., Chivers, D.P., 2010. Chemical ecology of predator-prey interactions in aquatic ecosystems: A review and prospectus. Can. J. Zool. 88, 698–724. 10.1139/Z10-029

Gall, B.G., Brodie, E.D., 2009. Behavioral avoidance of injured conspecific and predatory chemical stimuli by larvae of the aquatic caddisfly Hesperophylax occidentalis. Can. J. Zool. 87, 1009– 1015. 10.1139/Z09-091

Gillmann, S.M., Hering, D., Lorenz, A.W., 2023. Habitat development and species arrival drive succession of the benthic invertebrate community in restored urban streams. Environ. Sci. Eur. 35. 10.1186/s12302-023-00756-x

Golovatyuk, L. V., Prokin, A.A., Nazarova, L.B., Zinchenko, T.D., 2022. Biodiversity, distribution and production of macrozoobenthos communities in the saline Chernavka River (Lake Elton basin, South-West Russia). Limnology 23, 337–353. 10.1007/s10201-021-00692-w

Gordon, T.A.C., Neto-Cerejeira, J., Furey, P.C., O’Gorman, E.J., 2018. Changes in feeding selectivity of freshwater invertebrates across a natural thermal gradient. Curr. Zool. 64, 231–242. 10.1093/cz/zoy011

Haq, S., Kaushal, S.S., Duan, S., 2018. Episodic salinization and freshwater salinization syndrome mobilize base cations, carbon, and nutrients to streams across urban regions. Biogeochemistry 141, 463–486. 10.1007/s10533-018-0514-2

Hart, B.T., Bailey, P., Edwards, R., Hortle, K., James, K., McMahon, A., Meredith, C., Swadling, K., 1991. A review of the salt sensitivity of the Australian freshwater biota. Hydrobiologia 210, 105–144. 10.1007/BF00014327

Heiman, D.R., Knight, A.W., 1975. The influence of temperature on the bioenergetics of the carnivorous stonefly nymph, Acroneuria californica Banks (Plecoptera: Perlidae). Ecology 56, 105–116. 10.2307/1935303

Helfman, G.S., 1989. Behavioral Ecology and Sociobiology Threat-sensitive predator avoidance in damselfish-trumpetfish interactions. Behav Ecol Sociobiol 24, 47–58.

Herbert-Read, J.E., Logendran, D., Ward, A.J.W., 2010. Sensory ecology in a changing world: Salinity alters conspecific recognition in an amphidromous fish, Pseudomugil signifer. Behav. Ecol. Sociobiol. 64, 1107–1115. 10.1007/s00265-010-0925-0

Heuschele, J., Lode, T., Andersen, T., Titelman, J., 2020. The hidden dimension: Context-dependent expression of repeatable behavior in copepods. Environ. Toxicol. Chem. 39, 1017–1026. 10.1002/etc.4688

Hintz, W.D., Relyea, R.A., 2017. A salty landscape of fear: responses of fish and zooplankton to freshwater salinization and predatory stress. Oecologia 185, 147–156. 10.1007/s00442-017-3925-1

Hogg, I.D., Williams, D.D., 1996. Response of stream invertebrates to a global-warming thermal regime: An ecosystem-level manipulation. Ecology 77, 395–407. 10.2307/2265617

Howard, G.J., Webster, T.F., 2009. Generalized concentration addition: A method for examining mixtures containing partial agonists. J. Theor. Biol. 259, 469–477. 10.1016/j.jtbi.2009.03.030

Huber, E.D., Wilmoth, B., Hintz, L.L., Horvath, A.D., McKenna, J.R., Hintz, W.D., 2023. Freshwater salinization reduces vertical movement rate and abundance of Daphnia: Interactions with predatory stress. Environ. Pollut. 330. 10.1016/j.envpol.2023.121767

Huryn, A.D., Chivers, D.P., 1999. Contrasting behavioral responses by detritivorous and predatory mayflies to chemicals released by injured conspecifics and their predators. J. Chem. Ecol. 25, 2729–2740. 10.1023/A:1020851524335

Jack, J.D., Fang, W., Thorp, J.H., 2006. Vertical, lateral and longitudinal movement of zooplankton in a large river. Freshw. Biol. 51, 1646–1654. 10.1111/j.1365-2427.2006.01600.x

Jackson, M.C., Pawar, S., Woodward, G., 2021. The temporal dynamics of multiple stressor effects: From individuals to ecosystems. Trends Ecol. Evol. 36, 402–410. 10.1016/j.tree.2021.01.005

Jones, D.K., Mattes, B.M., Hintz, W.D., Schuler, M.S., Stoler, A.B., Lind, L.A., Cooper, R.O., Relyea, R.A., 2017. Investigation of road salts and biotic stressors on freshwater wetland communities. Environ. Pollut. 221, 159–167. 10.1016/j.envpol.2016.11.060

Kail, J., Palt, M., Lorenz, A., Hering, D., 2021. Woody buffer effects on water temperature: The role of spatial configuration and daily temperature fluctuations. Hydrol. Process. 35, 1–12. 10.1002/hyp.14008

Kasumyan, A.O., 2022. Fish as sources of kairomones–Chemical signals for aquatic animals. J. Ichthyol. 62, 289–315. 10.1134/S0032945222020084

Kleiber, C., Zeileis, A., 2008. Applied Econometrics with R, Applied Econometrics with R. Springer New York, New York. 10.1007/978-0-387-77318-6

Kottelat, M., Freyhof, J., 2007. Handbook of European freshwater fishes, Copeia. Publications Kottelat, Cornol and Freyhof, Berlin.

Lancaster, J., 1990. Predation and drift of lotic macroinvertebrates during colonization. Oecologia 85, 48–56. 10.1007/BF00317342

Lauridsen, T.L., Lodge, D.M., 1996. Avoidance by Daphnia magna of fish and macrophytes: Chemical cues and predator-mediated use of macrophyte habitat. Limnol. Oceanogr. 41, 794–798. 10.4319/lo.1996.41.4.0794

Lee, C.E., Petersen, C.H., 2003. Effects of developmental acclimation on adult salinity tolerance in the freshwater-invading copepod Eurytemora affinis. Physiol. Biochem. Zool. 76, 296–301. 10.1086/375433

Lenth, R. V., 2023. emmeans: Estimated marginal means, aka least-squares means.

Lima, S.L., Dill, L.M., 1990. Behavioral decisions made under the risk of predation: a review and prospectus. Can. J. Zool. 68, 619–640. 10.1139/z90-092

Löfgren, S., 2001. The chemical effects of deicing salt on soil and stream water of five catchments in southeast Sweden. Water. Air. Soil Pollut. 130, 863–868. 10.1023/A:1013895215558

Lubini, V., Knispel, S., Vincon, G., 2012. Fauna Helvetica: Plecoptera Identification.

Madge Pimentel, I., Baikova, D., Buchner, D., Burfeid Castellanos, A., David, G.M., Deep, A., Doliwa, A., Hadžiomerović, U., Mayombo, N.A.S., Prati, S., Spyra, M.A., Vermiert, A.M., Beisser, D., Dunthorn, M., Piggott, J.J., Sures, B., Tiegs, S.D., Leese, F., Beermann, A.J., 2024a. Assessing the response of an urban stream ecosystem to salinization under different flow regimes. Sci. Total Environ. 926, 171849. 10.1016/j.scitotenv.2024.171849

Madge Pimentel, I., Rehsen, P.M., Beermann, A.J., Leese, F., Piggott, J.J., Schmuck, S., 2024b. An automated modular heating solution for experimental flow-through stream mesocosm systems. Limnol. Oceanogr. Methods 22, 135–148. 10.1002/lom3.10596

Magbanua, F.S., Townsend, C.R., Hageman, K.J., Piggott, J.J., Matthaei, C.D., 2016. Individual and combined effects of fine sediment and glyphosate herbicide on invertebrate drift and insect emergence: A stream mesocosm experiment. Freshw. Sci. 35, 139–151. 10.1086/684363

Müller, K., 1954. Investigations on the organic drift in North Swedish streams. Rep. Inst. Freshw. Res. Drottningholm 35, 133–148.

Ou, M., Hamilton, T.J., Eom, J., Lyall, E.M., Gallup, J., Jiang, A., Lee, J., Close, D.A., Yun, S.S., Brauner, C.J., 2015. Responses of pink salmon to CO2-induced aquatic acidification. Nat. Clim. Chang. 5, 950–957. 10.1038/nclimate2694

Padisak, J., 1992. Seasonal Succession of Phytoplankton in a Large Shallow Lake (Balaton, Hungary)--A Dynamic Approach to Ecological Memory, Its Possible Role and Mechanisms. J. Ecol. 80, 217–230. 10.2307/2261008

Pasternak, A.F., Mikheev, V.N., Wanzenböck, J., 2006. How plankton copepods avoid fish predation: From individual responses to variations of the life cycle. J. Ichthyol. 46, 220–226. 10.1134/S0032945206110129

Peacor, S.D., Werner, E.E., 1997. Trait-mediated indirect interactions in a simple aquatic food web. Ecology 78, 1146–1156. 10.1890/0012-9658(1997)078[1146:TMIIIA]2.0.CO;2

Peckarsky, B.L., McIntosh, A.R., 1998. Fitness and community consequences of avoiding multiple predators. Oecologia 113, 565–576. 10.1007/s004420050410

Piggott, J.J., Townsend, C.R., Matthaei, C.D., 2015. Climate warming and agricultural stressors interact to determine stream macroinvertebrate community dynamics. Glob. Chang. Biol. 21, 1887–1906. 10.1111/gcb.12861

Pinder, L.C.V., 1986. Biology of freshwater Chironomidae. Annu. Rev. Entomol. Vol. 31 1–23. 10.1146/annurev.en.31.010186.000245

Poff, N.L.R., DeCino, R.D., Ward, J. V., 1991. Size-dependent drift responses of mayflies to experimental hydrologic variation: active predator avoidance or passive hydrodynamic displacement? Oecologia 88, 577–586. 10.1007/BF00317723

Quinn, J.M., Steele, G.L., Hickey, C.W., Vickers, M.L., 1994. Upper thermal tolerances of twelve New Zealand stream invertebrate species. New Zeal. J. Mar. Freshw. Res. 28, 391–397. 10.1080/00288330.1994.9516629

R Core Team, 2021. R: a language and environment for statistical computing. Vienna: R Foundation for Statistical Computing.

Reustle, J.W., Smee, D.L., 2020. Turbidity and salinity influence trophic cascades on oyster reefs through modification of sensory performance and facilitation of different predator types. Mar. Ecol. Prog. Ser. 639, 127–136. 10.3354/meps13283

Ross, E., Behringer, D., 2019. Changes in temperature, pH, and salinity affect the sheltering responses of Caribbean spiny lobsters to chemosensory cues. Sci. Rep. 9, 1–11. 10.1038/s41598-019-40832-y

Schaefer, M., 2018. Brohmer – Fauna von Deutschland. Quelle & Meyer Verlag GmbH & Co.

Schäfer, R.B., Piggott, J.J., 2018. Advancing understanding and prediction in multiple stressor research through a mechanistic basis for null models. Glob. Chang. Biol. 24, 1817–1826. 10.1111/gcb.14073

Schäffer, M., Winkelmann, C., Hellmann, C., Benndorf, J., 2013. Reduced drift activity of two benthic invertebrate species is mediated by infochemicals of benthic fish. Aquat. Ecol. 47, 99–107. 10.1007/s10452-013-9428-1

Schröder, M., Sondermann, M., Sures, B., Hering, D., 2015. Effects of salinity gradients on benthic invertebrate and diatom communities in a German lowland river. Ecol. Indic. 57, 236–248. 10.1016/j.ecolind.2015.04.038

Shtull-Trauring, E., Cohen, A., Ben-Hur, M., Tanny, J., Bernstein, N., 2020. Reducing salinity of treated waste water with large scale desalination. Water Res. 186, 116322. 10.1016/j.watres.2020.116322

Sih, A., 1980. Optimal Behavior: Can Foragers Balance Two Conflicting Demands? Science (80-.). 210, 1041–1043. 10.1126/science.210.4473.1041

Stabell, O.B., Ogbebo, F., Primicerio, R., 2003. Inducible defences in Daphnia depend on latent alarm signals from conspecific prey activated in predators. Chem. Senses 28, 141–153. 10.1093/chemse/28.2.141

Suarez, R.K., 2012. Energy and metabolism. Compr. Physiol. 2, 2527–2540. 10.1002/cphy.c110009

Sundermann, A., Lohse, S., 2004. Bestimmungsschlüssel für die aquatischen Zweiflügler (Diptera) in Anleh-nung an die Operationelle Taxaliste für Fließgewässer in Deutschland.

Tollrian, R., Duggen, S., Weiss, L.C., Laforsch, C., Kopp, M., 2015. Density-dependent adjustment of inducible defenses. Sci. Rep. 5, 1–9. 10.1038/srep12736

Tollrian, R., Harvell, C.D., 1999. The Ecology and Evolution of Inducible Defenses, The Quarterly Review of Biology. Princeton University Press, Princeton. 10.1515/9780691228198

Trotter, B., Ramsperger, A.F.R.M., Raab, P., Haberstroh, J., Laforsch, C., 2019. Plastic waste interferes with chemical communication in aquatic ecosystems. Sci. Rep. 9, 1–8. 10.1038/s41598-019-41677-1

Van Donk, E., Peacor, S., Grosser, K., De Senerpont Domis, L.N., Lürling, M., 2015. Pharmaceuticals may disrupt natural chemical information flows and species interactions in aquatic systems: Ideas and perspectives on a hidden Global Change, in: Reviews of Environmental Contamination and Toxicology. pp. 91–105. 10.1007/398_2015_5002

Vannote, R.L., Sweeney, B.W., 1980. Geographic analysis of thermal equilibria: A conceptual model for evaluating the effect of natural and modified thermal regimes on aquatic insect communities. Am. Nat. 115, 667–695. 10.1086/283591

Wagenhoff, A., Townsend, C.R., Matthaei, C.D., 2012. Macroinvertebrate responses along broad stressor gradients of deposited fine sediment and dissolved nutrients: A stream mesocosm experiment. J. Appl. Ecol. 49, 892–902. 10.1111/j.1365-2664.2012.02162.x

Waringer, J., Graf, W., 1997. Atlas der österreichischen Köcherfliegenlarven. Facultas Universitätsverlag, Wien.

Waters, T.F., 1972. The drift of stream insects. Annu. Rev. Entomol. 17, 253–272. 10.1146/annurev.en.17.010172.001345

Weiss, L.C., 2019. Sensory ecology of predator-induced phenotypic plasticity. Front. Behav. Neurosci. 12, 1–12. 10.3389/fnbeh.2018.00330

Weiss, L.C., Pötter, L., Steiger, A., Kruppert, S., Frost, U., Tollrian, R., 2018. Rising pCO2 in freshwater ecosystems has the potential to negatively affect predator-induced defenses in Daphnia. Curr. Biol. 28, 327–332. 10.1016/j.cub.2017.12.022

Weiss, L.C., Tollrian, R., 2018. Predator-induced defenses in crustacea, in: Life Histories. Oxford University PressNew York, pp. 303–322. 10.1093/oso/9780190620271.003.0012

Wickham, H., Averick, M., Bryan, J., Chang, W., McGowan, L., François, R., Grolemund, G., Hayes, A., Henry, L., Hester, J., Kuhn, M., Pedersen, T., Miller, E., Bache, S., Müller, K., Ooms, J., Robinson, D., Seidel, D., Spinu, V., Takahashi, K., Vaughan, D., Wilke, C., Woo, K., Yutani, H., 2019. Welcome to the Tidyverse. J. Open Source Softw. 4, 1686. 10.21105/joss.01686

Willems, D.J., Kumar, A., Nugegoda, D., 2022. The acute toxicity of salinity in onshore unconventional gas waters to freshwater invertebrates in receiving environments: A systematic review. Environ. Toxicol. Chem. 41, 2928–2949. 10.1002/etc.5492

Williams, D.D., Moore, K.A., 1982. The effect of environmental factors on the activity of Gammarus pseudolimnaeus (Amphipoda). Hydrobiologia 96, 137–147. 10.1007/BF02185429

Williams, L.R., Taylor, C.M., Warren, M.L., 2003. Influence of fish predation on assemblage structure of macroinvertebrates in an intermittent stream. Trans. Am. Fish. Soc. 132, 120–130. 10.1577/1548-8659(2003)132<0120:iofpoa>2.0.co;2

Wisenden, B.D., Chivers, D.P., Smith, R.J.F., 1997. Learned recognition of predation risk by Enallagma damselfly larvae (Odonata, Zygoptera) on the basis of chemical cues. J. Chem. Ecol. 23, 137–151. 10.1023/B:JOEC.0000006350.66424.3d

Wood, P.J., Dykes, A.P., 2002. The use of salt dilution gauging techniques: Ecological considerations and insights. Water Res. 36, 3054–3062. 10.1016/S0043-1354(01)00519-X

Woodward, G., Perkins, D.M., Brown, L.E., 2010. Climate change and freshwater ecosystems: Impacts across multiple levels of organization. Philos. Trans. R. Soc. B Biol. Sci. 365, 2093– 2106. 10.1098/rstb.2010.0055

Zaret, T.M., Suffern, J.S., 1976. Vertical migration in zooplankton. Limnol. Oceanogr. 21, 804–813.

Zinchenko, T.D., Golovatyuk, L. V., 2013. Salinity tolerance of macroinvertebrates in stream waters (review). Arid Ecosyst. 3, 113–121. 10.1134/S2079096113030116

