## Supplementary fig. 1-2 and tab. 1-3 for "Salinisation and warming disrupt predator-induced drift behaviour in aquatic predator-prey interactions"

### Supplement

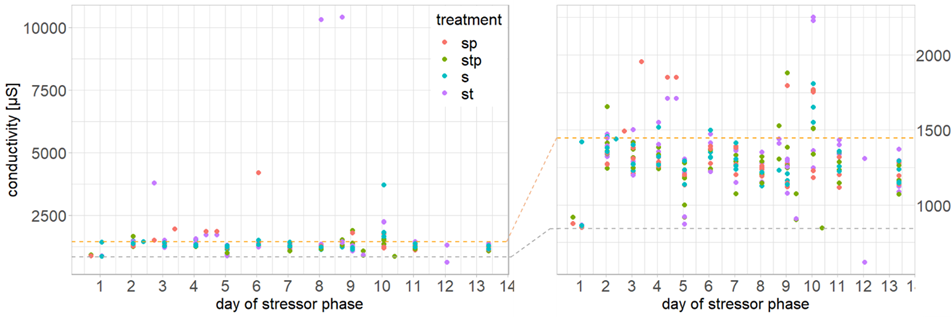

Supplementary fig. 1: Conductivity fluctuations of the stressor mesocosms during the 14-day stressor phase of the *ExStream* system. Mesocosms undergoing the treatment salinity (s), salinity and temperature (st), salinity and predator exposure (sp) and salinity, temperature, and predator exposure (stp) were continuously supplied with a diluted NaCl solution (refined salt tablets, Claramat, > 99.9 % NaCl, >350 mg/L NaCl) via an irrigation line using a pressure-compensated dripper system. The conductivity rose from an average of 840.63 µS ± 94.20 SD (grey dash line) under ambient conditions to an average of 1446.06 µS ± 912.39 SD (orange dash line).

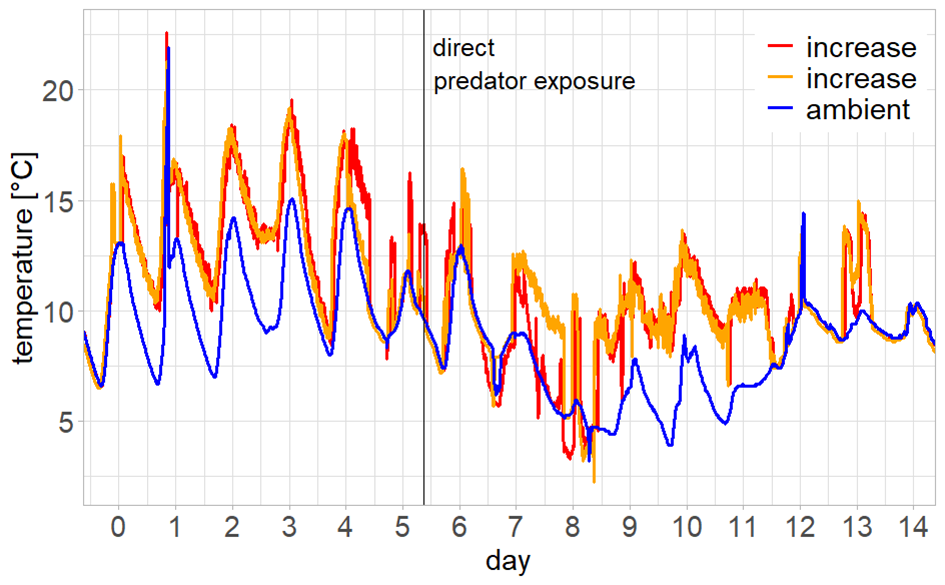

Supplementary fig. 2: Temperature fluctuations during the stressor phase of 14 days. The water temperature of two header tanks (orange and red line) of the *ExStream* system, were heated to + 4.5 °C (± 0.2 °C) relative to the ambient water temperature (blue line). Four header tanks each supplied 16 mesocosm with water.

Supplementary tab. 1: Summary of post hoc comparisons among groups results (emmeans). The number of drifting invertebrates were fitted with a generalised linear model (glm). Salinity, temperature, indirect and direct predation exposure was examined under different conditions [emmeans(model, ~predation*temperature*salinity) %>% pairs(simple = "each", adjust = "fdr")]. The emmeans results were averaged over the levels of day. Predator-specific cues were introduced continuously into the system for indirect predator exposure. On the fifth day, predator exposure was augmented by introducing bullheads into the mesocosms. Bold font denotes significant (< 0.05) p-values.

| **taxa** | **condition** | **contrast** | **estimate** | **SE** | **df** | **z.ratio** | **p.value** |
| --- | --- | --- | --- | --- | --- | --- | --- |
| **overall** | predation |  |  |  |  |  |  |
|  | control | no - indirect | -26.480 | 10.17 | Inf | -2.604 | **0.014** |
|  |  | no - direct | -32.700 | 8.42 | Inf | -3.885 | **< 0.001** |
|  |  | indirect - direct | -6.220 | 12.12 | Inf | -0.513 | 0.608 |
|  | increased salinity | no - indirect | 5.920 | 8.14 | Inf | 0.727 | 0.681 |
|  |  | no - direct | -3.510 | 8.54 | Inf | -0.411 | 0.681 |
|  |  | indirect - direct | -9.430 | 10.27 | Inf | -0.919 | 0.681 |
|  | increased temperature | no - indirect | 1.930 | 9.03 | Inf | 0.213 | 0.831 |
|  |  | no - direct | -15.920 | 7.64 | Inf | -2.083 | 0.112 |
|  |  | indirect - direct | -17.850 | 10.62 | Inf | -1.680 | 0.139 |
|  | increased salinity and temperature | no - indirect | -9.350 | 9.15 | Inf | -1.023 | 0.306 |
|  |  | no - direct | -33.580 | 8.06 | Inf | -4.167 | **< 0.001** |
|  |  | indirect - direct | -24.220 | 11.28 | Inf | -2.148 | **0.048** |
|  | temperature |  |  |  |  |  |  |
|  | control | ambient - increased | -2.780 | 5.94 | Inf | -0.468 | 0.640 |
|  | IP (indirect predation) | ambient - increased | 25.620 | 11.51 | Inf | 2.227 | **0.026** |
|  | DP (direct predation) | ambient - increased | 14.000 | 9.50 | Inf | 1.474 | 0.141 |
|  | increased salinity | ambient - increased | 10.390 | 5.84 | Inf | 1.779 | 0.075 |
|  | IP and salinity | ambient - increased | -4.880 | 10.17 | Inf | -0.480 | 0.631 |
|  | DP and salinity | ambient - increased | -19.680 | 10.01 | Inf | -1.965 | **0.050** |
|  | salinity |  |  |  |  |  |  |
|  | control | ambient - increased | -7.010 | 6.11 | Inf | -1.148 | 0.251 |
|  | IP (indirect predation) | ambient - increased | 25.380 | 11.00 | Inf | 2.307 | **0.021** |
|  | DP (direct predation) | ambient - increased | 22.180 | 10.12 | Inf | 2.191 | **0.029** |
|  | increased temperature | ambient - increased | 6.160 | 5.66 | Inf | 1.088 | 0.277 |
|  | IP and temperature | ambient - increased | -5.120 | 10.71 | Inf | -0.479 | 0.632 |
|  | DP and temperature | ambient - increased | -11.500 | 9.39 | Inf | -1.225 | 0.221 |
| **chironomid larvae** | predation |  |  |  |  |  |  |
|  | control | no - indirect | -7.517 | 2.75 | Inf | -2.729 | **0.010** |
|  |  | no - direct | -5.877 | 1.91 | Inf | -3.080 | **0.006** |
|  |  | indirect - direct | 1.641 | 3.15 | Inf | 0.521 | 0.602 |
|  | increased salinity | no - indirect | 3.775 | 1.94 | Inf | 1.945 | 0.155 |
|  |  | no - direct | 0.563 | 2.05 | Inf | 0.274 | 0.784 |
|  |  | indirect - direct | -3.212 | 2.35 | Inf | -1.369 | 0.257 |
|  | increased temperature | no - indirect | 0.732 | 2.31 | Inf | 0.316 | 0.752 |
|  |  | no - direct | -3.252 | 1.76 | Inf | -1.853 | 0.192 |
|  |  | indirect - direct | -3.984 | 2.65 | Inf | -1.501 | 0.200 |
|  | increased salinity and temperature | no - indirect | 0.536 | 2.24 | Inf | 0.240 | 0.811 |
|  |  | no - direct | -5.823 | 1.83 | Inf | -3.177 | **0.005** |
|  |  | indirect - direct | -6.359 | 2.70 | Inf | -2.353 | **0.028** |
|  | temperature |  |  |  |  |  |  |
|  | control | ambient - increased | -0.624 | 1.32 | Inf | -0.471 | 0.637 |
|  | IP (indirect predation) | ambient - increased | 7.625 | 3.13 | Inf | 2.434 | **0.015** |
|  | DP (direct predation) | ambient - increased | 2.000 | 2.20 | Inf | 0.908 | 0.364 |
|  | increased salinity | ambient - increased | 3.437 | 1.44 | Inf | 2.387 | **0.017** |
|  | IP and salinity | ambient - increased | 0.198 | 2.44 | Inf | 0.081 | 0.936 |
|  | DP and salinity | ambient - increased | -2.950 | 2.32 | Inf | -1.272 | 0.203 |
|  | salinity |  |  |  |  |  |  |
|  | control | ambient - increased | -3.115 | 1.49 | Inf | -2.092 | **0.036** |
|  | IP (indirect predation) | ambient - increased | 8.177 | 2.91 | Inf | 2.809 | **0.005** |
|  | DP (direct predation) | ambient - increased | 3.325 | 2.34 | Inf | 1.419 | 0.156 |
|  | increased temperature | ambient - increased | 0.946 | 1.27 | Inf | 0.747 | 0.455 |
|  | IP and temperature | ambient - increased | 0.750 | 2.70 | Inf | 0.277 | 0.782 |
|  | DP and temperature | ambient - increased | -1.625 | 2.18 | Inf | -0.747 | 0.455 |
| **micro-crustaceans** | predation |  |  |  |  |  |  |
|  | control | no - indirect | -14.868 | 6.28 | Inf | -2.368 | **0.027** |
|  |  | no - direct | -16.620 | 5.29 | Inf | -3.139 | **0.005** |
|  |  | indirect - direct | -1.752 | 7.51 | Inf | -0.233 | 0.816 |
|  | increased salinity | no - indirect | 0.789 | 5.07 | Inf | 0.156 | 0.995 |
|  |  | no - direct | -0.033 | 5.09 | Inf | -0.006 | 0.995 |
|  |  | indirect - direct | -0.822 | 6.26 | Inf | -0.131 | 0.995 |
|  | increased temperature | no - indirect | 2.089 | 5.53 | Inf | 0.378 | 0.706 |
|  |  | no - direct | -7.163 | 4.86 | Inf | -1.473 | 0.236 |
|  |  | indirect - direct | -9.252 | 6.54 | Inf | -1.415 | 0.236 |
|  | increased salinity and temperature | no - indirect | -9.552 | 5.72 | Inf | -1.669 | 0.143 |
|  |  | no - direct | -17.554 | 4.94 | Inf | -3.553 | **0.001** |
|  |  | indirect - direct | -8.002 | 7.03 | Inf | -1.138 | 0.255 |
|  | temperature |  |  |  |  |  |  |
|  | control | ambient - increased | -2.210 | 3.81 | Inf | -0.58 | 0.562 |
|  | IP (indirect predation) | ambient - increased | 14.750 | 7.04 | Inf | 2.096 | **0.036** |
|  | DP (direct predation) | ambient - increased | 7.250 | 5.96 | Inf | 1.217 | 0.224 |
|  | increased salinity | ambient - increased | 5.650 | 3.56 | Inf | 1.586 | 0.113 |
|  | IP and salinity | ambient - increased | -4.700 | 6.41 | Inf | -0.733 | 0.464 |
|  | DP and salinity | ambient - increased | -11.88 | 6.02 | Inf | -1.973 | 0.049 |
|  | salinity |  |  |  |  |  |  |
|  | control | ambient - increased | -2.210 | 3.81 | Inf | -0.581 | 0.561 |
|  | IP (indirect predation) | ambient - increased | 13.450 | 6.81 | Inf | 1.973 | 0.049 |
|  | DP (direct predation) | ambient - increased | 14.380 | 6.14 | Inf | 2.339 | **0.019** |
|  | increased temperature | ambient - increased | 5.640 | 3.56 | Inf | 1.584 | 0.113 |
|  | IP and temperature | ambient - increased | -6.000 | 6.64 | Inf | -0.903 | 0.366 |
|  | DP and temperature | ambient - increased | -4.750 | 5.83 | Inf | -0.815 | 0.415 |

Supplementary tab. 2: Summary of post hoc comparisons among groups results (emmeans). The number of invertebrates in the leaf litter were fitted with a generalised linear model (glm). Predation exposure, temperature and salinity was examined under different conditions [emmeans(model, ~predation*temperature*salinity) %>% pairs(simple = "each", adjust = "fdr")]. Two-way interactions were found to affect the Tanytarsini larvae, Chironomidae pupae and gammarids. Three-way interaction affected Trichoptera > 3 mm and Harpacticoida [emmeans(model, specs = "", by = c("", “”)) %>% pairs(simple = "each", adjust = "fdr")]. The results for Tanytarsini larvae and Chironomidae pupae were averaged over the level temperature and over the level salinity for gammarids. Predator-specific cues were introduced continuously into the system for indirect predator exposure. On the fifth day, predator exposure was augmented by introducing bullheads into the mesocosms. Bold and italic font denotes tendential (< 0.05) and bold font denotes significant (< 0.1) p-values.

| **group** | **condition** | **contrast** | **estimate** | **SE** | **df** | **z.ratio** | **p.value** |
| --- | --- | --- | --- | --- | --- | --- | --- |
| **Tanytarsini larvae** | no predation | ambient - increased | -12.46 | 3.13 | Inf | -3.974 | **< 0.001** |
|  | predation | ambient - increased | -2.12 | 3.11 | Inf | -0.684 | 0.494 |
|  | ambient salinity | no - predation | -2.75 | 2.53 | Inf | -1.087 | 0.277 |
|  | increased salinity | no - predation | 7.58 | 3.62 | Inf | 2.096 | **0.036** |
| **Chironomidae pupae** | no predation | ambient - increased | -5.12 | 1.64 | Inf | -3.123 | **0.002** |
|  | predation | ambient - increased | -0.75 | 1.65 | Inf | -0.455 | 0.649 |
|  | ambient salinity | no - predation | -1.38 | 1.40 | Inf | -0.984 | 0.325 |
|  | increased salinity | no - predation | 3.00 | 1.86 | Inf | 1.613 | 0.107 |
| **gammarids** | no predation | ambient - increased | 9.12 | 8.52 | Inf | 1.071 | 0.284 |
|  | predation | ambient - increased | -18.25 | 9.00 | Inf | -2.028 | **0.043** |
|  | ambient temperature | no - predation | 11.08 | 9.06 | Inf | 1.297 | 0.195 |
|  | increased temperature | no - predation | -15.60 | 8.46 | Inf | -1.847 | ***0.065*** |
| **Harpacticoida** | control | no - predation | -12.25 | 20.00 | Inf | -0.613 | 0.540 |
|  | increased salinity | no - predation | 24.00 | 28.70 | Inf | 0.837 | 0.403 |
|  | increased temperature | no - predation | 39.50 | 21.20 | Inf | 1.859 | ***0.063*** |
|  | increased salinity and temperature | no - predation | -7.25 | 19.90 | Inf | -0.364 | 0.716 |
| *temperature* | control | ambient - increased | -32.80 | 21.80 | Inf | -1.500 | 0.134 |
|  | predation | ambient - increased | 19.00 | 19.30 | Inf | 0.982 | 0.326 |
|  | increased salinity | ambient - increased | 30.80 | 24.20 | Inf | 1.270 | 0.204 |
|  | predation and increased salinity | ambient - increased | -0.50 | 25.20 | Inf | -0.020 | 0.984 |
| *salinity* | control | ambient - increased | -33.00 | 24.00 | Inf | -1.373 | 0.170 |
|  | predation | ambient - increased | 3.25 | 25.40 | Inf | 0.128 | 0.898 |
|  | increased temperature | ambient - increased | 30.50 | 22.00 | Inf | 1.385 | 0.166 |
|  | predation and increased temperature | ambient - increased | -16.25 | 19.10 | Inf | -0.852 | 0.394 |
| **Trichoptera**  **> 3 mm** | control | no - predation | -1.75 | 1.89 | Inf | -0.927 | 0.354 |
|  | increased salinity | no - predation | -7.17 | 2.68 | Inf | -2.672 | **0.008** |
|  | increased temperature | no - predation | -2.50 | 1.66 | Inf | -1.508 | 0.132 |
|  | increased salinity and temperature | no - predation | 2.50 | 1.90 | Inf | 1.313 | 0.189 |
| *temperature* | control | ambient - increased | 2.00 | 1.62 | Inf | 1.234 | 0.217 |
|  | predation | ambient - increased | 1.25 | 1.92 | Inf | 0.651 | 0.515 |
|  | increased salinity | ambient - increased | -4.17 | 1.89 | Inf | -2.205 | **0.027** |
|  | predation and increased salinity | ambient - increased | 5.50 | 2.69 | Inf | 2.043 | **0.041** |
| *salinity* | control | ambient - increased | 1.92 | 1.73 | Inf | 1.105 | 0.269 |
|  | predation | ambient - increased | -3.50 | 2.78 | Inf | -1.257 | 0.209 |
|  | increased temperature | ambient - increased | -4.25 | 1.79 | Inf | -2.380 | **0.017** |
|  | predation and increased temperature | ambient - increased | 0.75 | 1.79 | Inf | 0.420 | 0.674 |

Supplementary tab. 3: Summary of post hoc comparisons among groups results (emmeans). The invertebrate numbers of the community were fitted with a generalised linear model (glm). Predation exposure, temperature and salinity was examined under different conditions [emmeans(model, ~predation*temperature*salinity) %>% pairs(simple = "each", adjust = "fdr")]. Two-way interactions affected the Ceratopogonidae larvae and Trichoptera over 3 mm. Three-way interaction affected the Chironomidae pupae. The results for Ceratopogonidae larvae and Trichoptera over 3 mm were averaged over the level salinity. Chemical cues were introduced continuously into the system for indirect predation. On the fifth day, predator exposure was augmented by introducing bullheads into the mesocosms for direct predation. Bold font denotes significant (< 0.05) p-values.

| **group** | **condition** | **contrast** | **estimate** | **SE** | **df** | **z.ratio** | **p.value** |
| --- | --- | --- | --- | --- | --- | --- | --- |
| **Chironomidae**  **pupae** | control | no - predation | -3.50 | 2.65 | Inf | -1.323 | 0.186 |
|  | increased salinity | no - predation | 5.75 | 3.15 | Inf | 1.824 | **0.068** |
|  | increased temperature | no - predation | 1.83 | 3.44 | Inf | 0.532 | 0.595 |
|  | increased salinity and temperature | no - predation | -8.25 | 3.23 | Inf | -2.554 | **0.011** |
| *temperature* | control | ambient - increased | -10.50 | 2.96 | Inf | -3.550 | **< 0.001** |
|  | predation | ambient - increased | -1.25 | 2.86 | Inf | -0.437 | 0.662 |
|  | increased salinity | ambient - increased | -1.42 | 3.05 | Inf | -0.465 | 0.642 |
|  | predation and increased salinity | ambient - increased | -11.50 | 3.61 | Inf | -3.190 | **0.001** |
| *salinity* | control | ambient - increased | -3.08 | 2.86 | Inf | -1.078 | 0.281 |
|  | predation | ambient - increased | 2.25 | 3.27 | Inf | 0.688 | 0.491 |
|  | increased temperature | ambient - increased | 6.00 | 3.14 | Inf | 1.909 | ***0.056*** |
|  | predation and increased temperature | ambient - increased | -8.00 | 3.24 | Inf | -2.469 | **0.014** |
| **Cerato-pogonidae**  **larvae** | no predation | ambient - increased | 1.50 | 4.61 | Inf | 0.326 | 0.745 |
|  | predation | ambient - increased | 18.10 | 5.14 | Inf | 3.529 | **< 0.001** |
|  | ambient temperature | no - predation | -8.00 | 5.59 | Inf | -1.432 | 0.152 |
|  | increased temperature | no - predation | 8.62 | 4.05 | Inf | 2.130 | **0.033** |
| **Trichoptera**  **> 3 mm** | no predation | ambient - increased | 0.46 | 3.44 | Inf | 0.133 | 0.894 |
|  | predation | ambient - increased | 10.75 | 4.39 | Inf | 2.448 | **0.014** |
|  | ambient temperature | no - predation | -9.29 | 4.55 | Inf | -2.042 | **0.041** |
|  | increased temperature | no - predation | 1.00 | 3.22 | Inf | 0.310 | 0.756 |
